# Cardiolipin constrains lipid unsaturation during anaerobic adaptation in *Escherichia coli*

**DOI:** 10.64898/2026.03.04.709520

**Authors:** Milica Denic, Manon Delprat, Leon Espinosa, Aude E. Barani, Farida Seduk, Yoshiki Yamaryo-Botté, Cyrille Botté, Stéphane Audebert, Luc Camoin, Axel Magalon

**Affiliations:** Aix Marseille Univ., CNRS, Laboratoire de Chimie Bactérienne (UMR7283), IMM, Marseille, France; Aix Marseille Univ., Université de Toulon, CNRS, IRD, Mediterranean Institute of Oceanography (MIO, UMR110), 13288 Marseille, France; Apicolipid Team & Gemeli Platform, Institute for Advanced Biosciences, CNRS UMR5309, INSERM U1209, Université Grenoble Alpes, Grenoble, France; Aix-Marseille Université, Inserm, CNRS, Marseille, Institut Paoli-Calmettes, Centre de Recherche en Cancérologie de Marseille (CRCM), Marseille Proteomics, Marseille, France

## Abstract

Membrane lipids play a crucial role in cellular adaptation; though their specific functions in bacterial adaptation to oxygen limitation are not yet fully elucidated. Cardiolipin (CL), a signature phospholipid in energy-transducing membranes and a key organizer of mitochondrial respiration, has an unclear role in bacterial anaerobic physiology. Here, genetics, quantitative lipidomics and proteomics, enzymology, and fluorescence imaging were combined to explore how CL facilitates hypoxia adaptation in *Escherichia coli*. ClsA emerged as the predominant CL synthase under anaerobic conditions, and CL deficiency selectively impaired nitrate-dependent growth, while fermentation and fumarate respiration remained largely unaffected. CL depletion triggered extensive lipidome rewiring during the aerobic-to-anaerobic transition, including increased levels of phosphatidylglycerol, phosphatidic acid, and diacylglycerol, along with a broad enrichment of more unsaturated lipid species. In parallel, proteome remodeling linked CL loss to reduced abundance of proteins involved in anoxic respiration and nitrosative stress management, alongside the induction of membrane stress responses. Interestingly, CL deficiency did not markedly affect cell morphology, nor the spatial distribution or stability of respiratory complexes. This suggests a specialized role in optimizing membrane protein function rather than providing generic structural support. Wild-type lipidome profiling further showed that anaerobiosis induces a shift of lipid species toward higher unsaturation, with only modest class-level changes. Collectively, these results connect lipid remodeling to functional outcomes *in vivo*, offering mechanistic insights into how bacteria adapt their membranes to maintain energy conservation in fluctuating environments.

## Introduction

Biological membranes function as dynamic platforms where lipid composition, protein abundance, and supramolecular organization are continuously adjusted to meet environmental challenges. This adaptability is particularly crucial in energy-transducing membranes, where electron transfer enzymes must maintain both catalytic efficiency and structural stability within a chemically and physically heterogeneous bilayer. A recurring principle observed across bacteria and mitochondria is that specific “signature” lipids can exert significant control over bioenergetic performance by shaping local membrane properties and stabilizing membrane protein assemblies [1,2]. Cardiolipin (CL) serves as a prime example. Its dimeric, tetra-acyl structure and anionic headgroup confer upon CL a non-bilayer-prone character that can promote negative curvature and domain formation [3–5]. In parallel, biochemical and structural studies consistently link CL to respiratory and photosynthetic enzymes, supporting optimal activity, stability, oligomerization, and higher-order organization of the proteins [6–8]. Beyond energy conversion, CL displays a versatile role in eukaryotic cell as a signaling lipid whose interactions with various partners can initiate or fine-tune cellular responses, including pathways linked to organelle quality control and inflammation [9–12]. In prokaryotes, CL similarly regulates critical adaptive systems, such as the SaeRS two-component system in *Staphylococcus aureus*, which controls virulence factor expression, and osmoprotective transporters like ProP by directly interacting with their membrane-embedded domains, modulating their conformation and signaling[1,13]. Importantly, the chemical state of CL, including its acyl-chain composition and susceptibility to oxidation, can further influence its interaction landscape, highlighting how a lipid can simultaneously function as a structural determinant and a condition-responsive regulator [9,14,15]. The dynamic properties of CL are closely tied to its fatty acid (FA) composition and redox state. However, these features are not static, they are highly responsive to environmental conditions, particularly oxygen availability [16].

Given that CL species enriched in unsaturated acyl chains are particularly susceptible to oxidation, a process directly influenced by O₂ fluctuations, this raises a parallel issue in bacteria that remains poorly addressed: to what extent does oxygen availability shape not only CL abundance but also its FA composition in organisms that routinely experience oxygen fluctuations? In *Escherichia coli*, this question is particularly compelling. Although CL constitutes only a minor component of the total glycerophospholipid pool, it is produced by three dedicated synthases (ClsA, ClsB, and ClsC), implying strong selective pressure to sustain its biosynthesis [17]. Moreover, CL has been implicated as a bound cofactor of several *E. coli* respiratory complexes belonging both to aerobic and anaerobic electron transfer chains, supported by structural observations and by lipid-dependent stimulation of enzyme activity, including formate dehydrogenase N, succinate dehydrogenase, NADH dehydrogenase, and nitrate reductase A [6,18–22]. This observation aligns with the broader hypothesis that CL preferentially occupies conserved sites at subunit or oligomer interfaces, where it can act as a flexible amphipathic “bridge” [23]. However, most functional insights into CL biology in *E. coli* have been derived under aerobic laboratory conditions, leaving its importance during anaerobiosis and the transition from aerobic to anaerobic metabolism, largely unexplored.

Oxygen availability is a dominant ecological constraint, and the capacity to sustain growth in its absence is fundamental to microbial survival. For facultative anaerobes such as *E. coli*, anaerobiosis is not merely an alternative state but a coordinated physiological program that preserves energy conservation and cellular homeostasis while electron acceptors and respiratory modules are reconfigured [24]. This transition is anticipated to impose multiple demands on the membrane, which must adapt to changes in the composition and activity of respiratory complexes while maintaining an environment conducive to efficient electron transfer. Whether CL contributes to this remodeling by modulating membrane properties, stabilizing key enzymes, or shaping their spatial organization remains an open question of direct relevance to the ecological niches encountered by *E. coli* [25].

Here, this knowledge gap is addressed by combining genetics, quantitative multi-omics, and targeted functional assays across anaerobic growth and the aerobic-to-anaerobic transition. Anaerobic fitness profiling of single and triple *cls* mutants identified nitrate respiration as the condition most sensitive to CL loss. Lipidomics and proteomics during the nitrate-respiring transition were then integrated with mechanistic analyses of two electron-connected respiratory complexes to link CL to enzyme activity and energy conservation. Finally, cell morphology and membrane localization were assessed to probe higher-order organization. Together, this framework connects CL-dependent membrane remodeling to respiratory performance during oxygen limitation and moves from lipid–protein association toward causal understanding *in vivo*.

## Results

### Cardiolipin is dispensable for most anaerobic regimes but is required for nitrate-dependent growth

To dissect the contribution of the three cardiolipin synthases to anaerobic physiology in *E. coli*, in-frame deletion mutants lacking *clsA* (Δ*clsA*), *clsB* (Δ*clsB*) or *clsC* (Δ*clsC*) were generated, along with a triple mutant strain lacking all three genes (Δ*clsABC*). These strains were then tested under distinct anaerobic energy regimes: glucose fermentation, which is largely independent of membrane processes except for substrate uptake and product export, and anaerobic respiration using fumarate or nitrate as terminal electron acceptors, both prevalent in ecological niches [26]. Given that CL has been identified in the structures of key nitrate-respiratory proteins such as NarI and FdnI [19,21] and thought to stabilize these complexes, we anticipated that cardiolipin deficiency might preferentially compromise nitrate-dependent growth. To isolate the contribution of nitrate reductase A (NarGHI) recognized as the major actor to nitrate respiration in the intestine [27], we used a genetic background devoid of nitrate reductase Z and the periplasmic nitrate reductase.

Consistent with this hypothesis, deletion of *clsA* resulted in a significant growth defect in nitrate respiration conditions, a phenotype recapitulated in the Δ*clsABC* strain, whereas Δ*clsB* and Δ*clsC* grew comparably to the wild-type strain (Fig. 1A). In contrast, all strains grew well under fumarate respiration conditions or glucose fermentation (Fig. 1B, C), consistent with previous observations that CL deficiency does not impair anaerobic respiration with TMAO [28]. These data identify nitrate respiration as the anaerobic regime most sensitive to CL loss and point to ClsA as the dominant contributor under these conditions [29,30]. To exclude gross morphological defects as a confounding factor, cell shape was quantified across the three regimes (Figure S1). No measurable changes in rod morphology, width, or length distributions were detected in any mutant background. Notably, nitrate-respiring cells displayed increased cell-length heterogeneity irrespective of genotype, indicating that this variability is a property of nitrate respiration itself (Figure S1). Collectively, these results indicate that the growth defect is not attributable to gross shape alterations.

**Figure 1.**
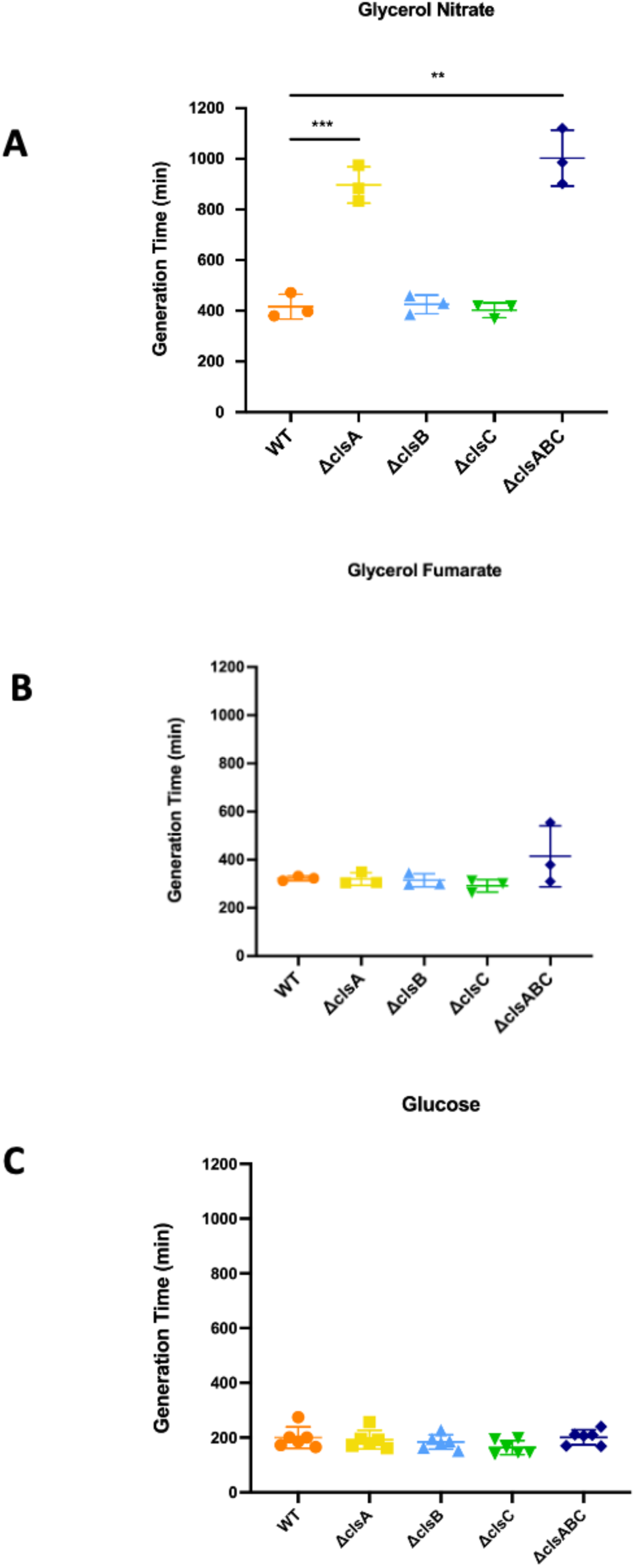
Inactivation of *clsA* and *clsABC* selectively impairs growth during nitrate respiration but not during fumarate respiration or glucose fermentation. Generation times of wild-type (WT), Δ*clsA*, Δ*clsB*, Δ*clsC*, and Δ*clsABC E. coli* strains were assessed under anaerobic conditions in defined medium. The strains were cultured with glycerol as the sole carbon source and nitrate **(A)**, or fumarate **(B)** as the sole electron acceptor in respiring conditions, or glucose as the sole carbon source in fermentative conditions **(C)**. Statistical analysis using the Unpaired t-test confirmed these differences as highly significant (*** *P* < 0.001; ** 0.001 ≤ *P* < 0.01).

### Cardiolipin is dispensable for complex stability but required for efficient energy conservation during nitrate respiration

As nitrate respiration in our strain background depends exclusively on nitrate reductase A (NarGHI) and CL has been shown to interact with key components of the formate-nitrate respiratory chain (including NarI and FdnI), our investigation focused on whether the observed growth phenotype in the Δ*clsABC* mutant, which is completely devoid of cardiolipin, arises from dysfunction within this electron transfer pathway at the level of its constituent complexes.

First, we measured formate dehydrogenase (Fdn) and nitrate reductase (Nar) enzyme activities in crude extract preparations. Fdn activity was measured *via* PMS-mediated reduction of DCPIP [29], an assay that selectively reports on periplasm-facing, membrane-inserted Fdn [30]. Nar activity was assayed using either reduced benzyl viologen (BV) or the quinol analog menadiol as electron donors [31,32]. While Fdn activity was indistinguishable between WT (0.9564 units) and Δ*clsABC* (1.041 units) (Fig.2A), Nar activity exhibited a donor-dependent phenotype: menadiol-supported rates were reduced by approximately 50% in Δc*lsABC* (4.47 units) compared to WT (9.25 units), indicating a significant impairment in quinol-linked activity (Fig. 2B), whereas BV-supported rates were unaffected (Fig. 2C). Since BV bypasses the membrane quinol-oxidation step, the relatively unchanged BV-driven Nar activity, along with normal Fdn activity, suggests that there is no major defect in the assembly or intrinsic stability of either complex. Instead, these findings highlight a CL-dependent step specifically linked to quinol utilization by NarGHI. This is consistent with prior evidence demonstrating that cardiolipin binding influences quinol substrate interactions [21].

**Figure 2.**
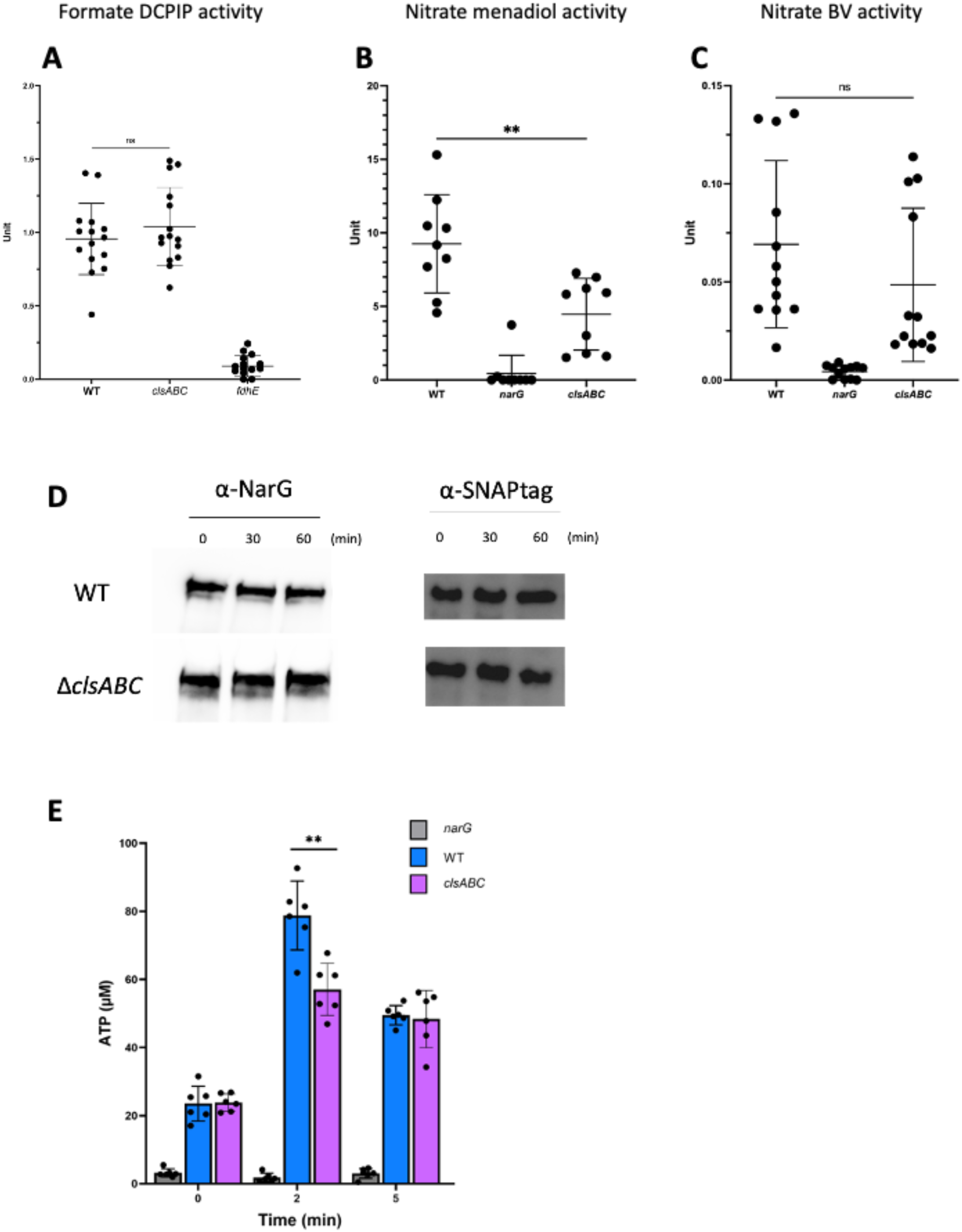
Functional analysis of Nar and Fdn activities and their impact on ATP production in WT and Δ*clsABC* strains. Fdn activity measured using DCPIP as an electron acceptor **(A)**, and Nar activity measured using menadiol **(B)** and reduced benzyl viologen (BV) **(C)** as electron donors, respectively. Western blot analysis of NarG and FdnI protein levels, fused with Halo and SNAP tags, respectively, in WT and Δ*clsABC* strains at 0, 30, and 60 minutes following a chloramphenicol chase **(D)**. ATP concentrations measured in nitrate-deficient (JCB4023), WT and Δ*clsABC* strains at 0, 2, and 5 minutes after initiating the reaction with 10 mM sodium formate and 100 mM potassium nitrate (KNO₃) **(E)**. Statistical analysis via **t-test** was performed to evaluate the significance of differences in enzyme activities and ATP levels between wild-type and Δ*clsABC* strains (ns *P* > 0.05; ** *P* < 0.01).

Next, we assessed the stability of each complexes using an orthogonal *in vivo* approach. WT and Δ*clsABC* strains expressing NarG::Halo_Tag_ and FdnI::SNAP_Tag_ were grown under nitrate respiration conditions and subjected to chloramphenicol chase, followed by immunoblotting (Fig. 2D). NarG::Halo_Tag_ and FdnI::SNAP_Tag_ remained stable and at comparable levels in both backgrounds, consistent with the high intrinsic stability of many respiratory complexes (Figure S3) [33]. Together, these data corroborate the enzymatic readouts and indicate that CL does not measurably affect the steady-state abundance or short-term stability of FdnGHI and NarGHI during nitrate respiration.

Finally, we investigated whether this enzymatic phenotype translated into altered energy conservation during formate-nitrate respiration, where the electrogenic activities of FdnGHI and NarGHI contribute to the formation of a transmembrane proton gradient. Resting cell suspensions displayed similar basal ATP pool concentrations in WT (23,52 µM) and Δ*clsABC* (22,82 µM) cells (Fig. 2E). Upon initiating respiration with formate and nitrate addition, WT cells showed a rapid ∼3-fold rise in ATP after 2 min (78,79 µM) whereas Δ*clsABC* exhibited a weaker increase (57,05 µM), with ATP levels converging by 5 min, consistent with the equilibration of cellular nucleotide pools.

Together, these results show that CL is not essential for Fdn activity and Nar/Fdn protein stability, but is required for efficient quinol-linked Nar activity and maximal ATP gain during nitrate respiration, underscoring a key role for CL in functional coupling within this electron transfer chain.

### Polar clustering of NarGHI and FdnGHI persists in the absence of cardiolipin during nitrate respiration

As previously described [34,35], formate dehydrogenase (FdnGHI) and nitrate reductase (NarGHI) complexes can assemble into large polar clusters and even interact under nitrate-respiring conditions. This spatial organization is proposed to enhance local quinone turnover, thereby supporting respiratory efficiency. Given the reduced functional coupling between FdnGHI and NarGHI in the Δ*clsABC* mutant, we sought to determine whether cardiolipin contributes to the membrane-scale positioning of these two electron-connected complexes during nitrate respiration. To address this, we performed fluorescence microscopy under nitrate-respiring conditions. Since GFP is inactive in anaerobic environments, we fused FdnI and NarG with HaloTag, allowing us to use organic dyes (HaloTag® TMR Ligand) for labeling and visualizing these proteins in the absence of oxygen [36,37].

For each strain, fluorescence intensity profiles were extracted along the cell medial axis (normalized from pole-to-pole) and aggregated across the population (Figure 3). In WT cells, the NarG signal displayed a pronounced bipolar enrichment, with sharp maxima at both poles and comparatively low intensity along the lateral membrane, consistent with polar clustering (Fig. 3A). In contrast, *clsABC* deficient strain exhibited a slightly altered profile: polar enrichment was less sharply defined, with elevated non-polar fluorescence compared to WT. A similar trend is observed when analyzing the spatial distribution of FdnI (Fig. 3B). Violin plots of the relative and longitudinal positions of both NarG and FdnI further supported these observations (Fig. 3C and 3D).

**Figure 3.**
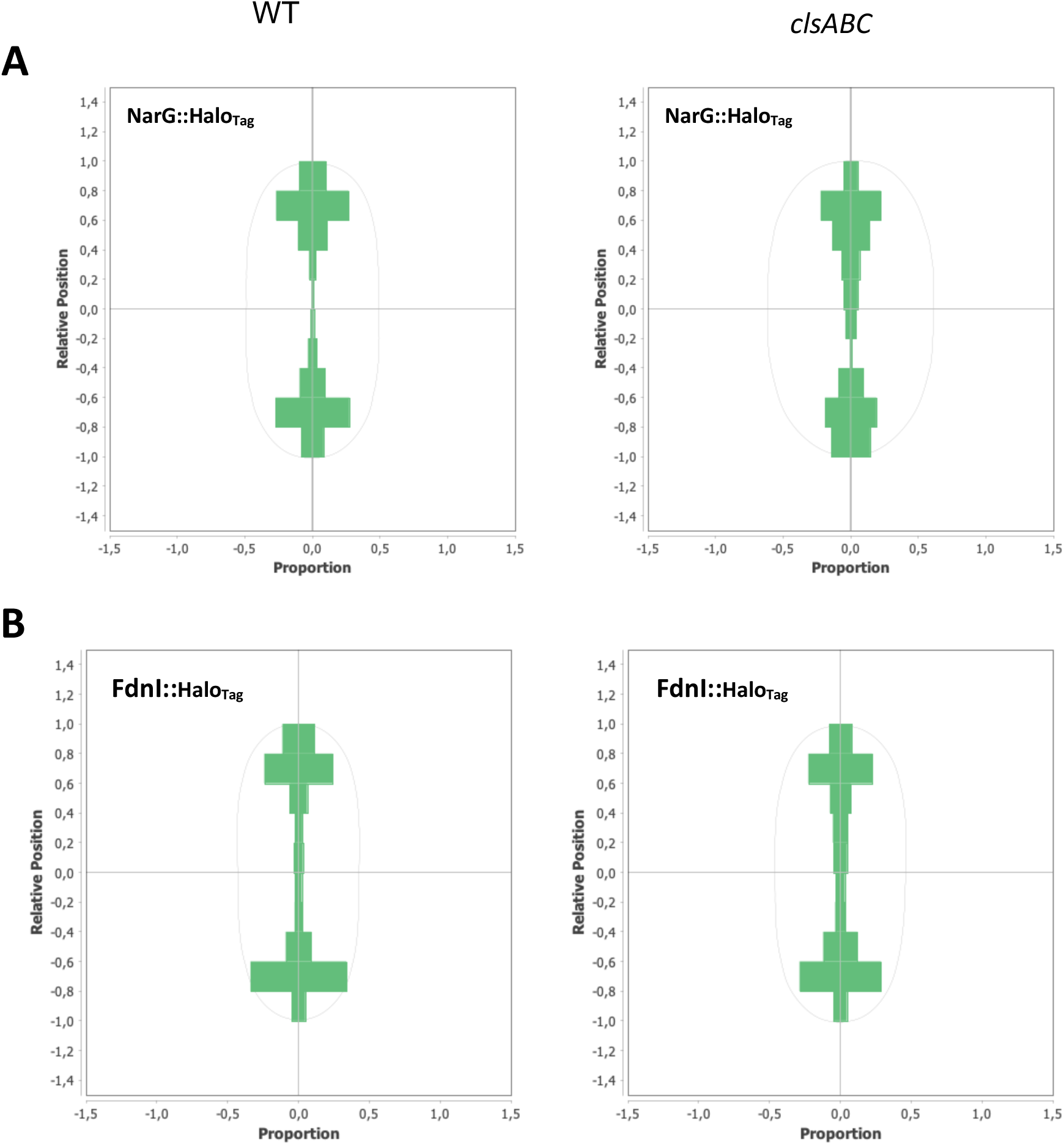
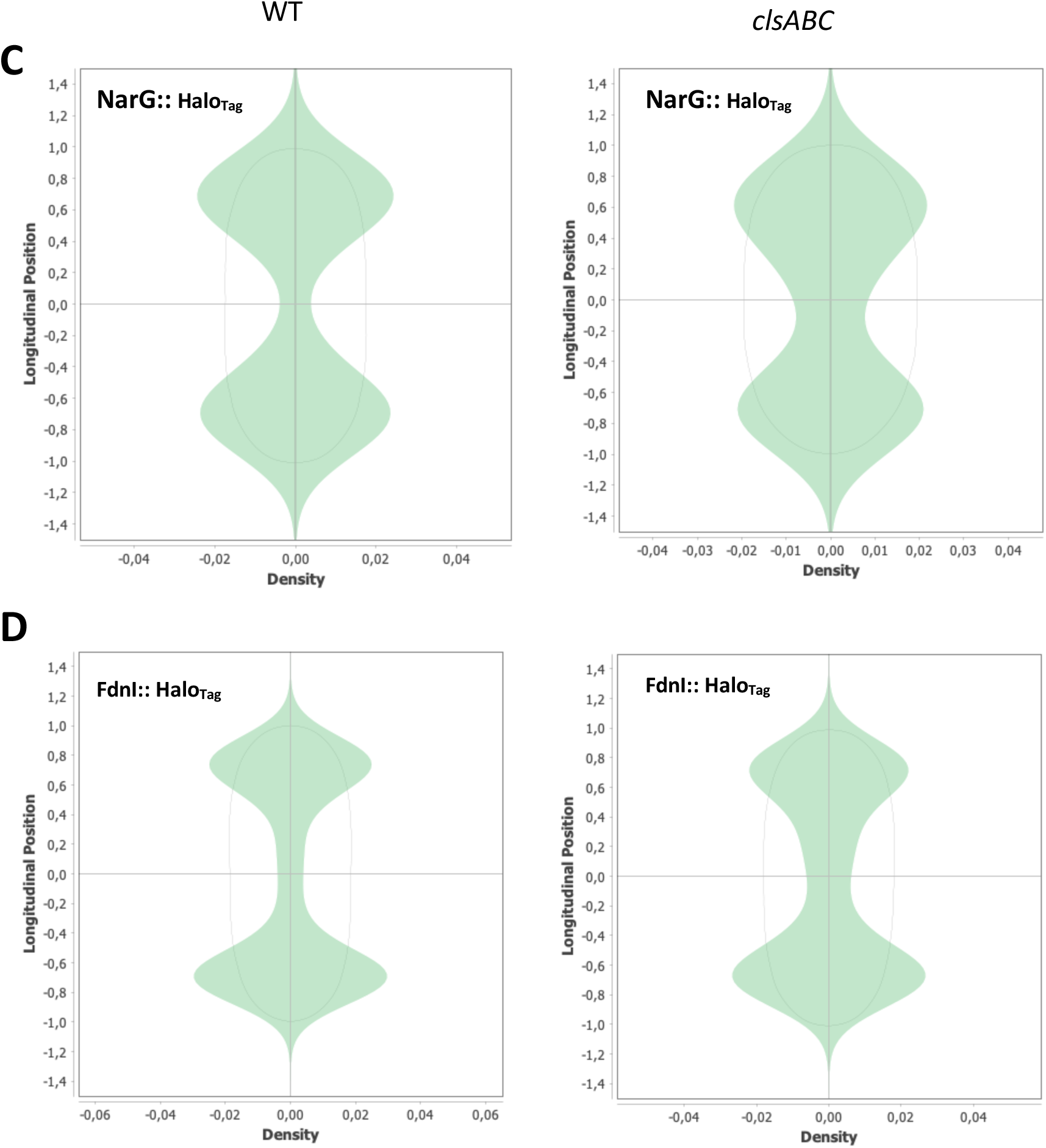
Polar localization of the NarGHI and FdnGHI complexes is maintained in a cardiolipin-deficient background during nitrate respiration. (**A, B**) Longitudinal vs. Positional Distribution: Scatter plots showing the spatial distribution of NarG::Halo**_Tag_** and FdnI::Halo**_Tag_** along the longitudinal axis of cells in WT and cardiolipin-deficient strains during nitrate respiration. The plots illustrate the positional localization patterns of the Halo-tagged complexes, with data points representing individual measurements across n=300 cells. (**C, D**) Relative Position Violin Plots: Violin plots depicting the density distribution of NarG::Halo**_Tag_** and FdnI::Halo**_Tag_** relative positions along the longitudinal axis in WT and cardiolipin-deficient strains. The distribution width and shape indicate the variability and concentration of protein localization. Data were collected from 300 cells across 3 independent experiments.

Notably, despite the altered distribution patterns, the ability of both FdnI and NarG to partition into polar clusters was not abolished in the Δ*clsABC* mutant. This suggests that polar clustering is robust to cardiolipin depletion and likely relies on additional determinants of membrane organization.

### Loss of cardiolipin triggers global lipidome rewiring and increased fatty-acid unsaturation during nitrate respiration

Given that CL is specifically required for nitrate-dependent anaerobic growth, we asked whether this phenotype reflects broader changes in membrane lipid homeostasis that could reshape the physicochemical environment supporting the nitrate-dependent respiration chain. We therefore focused our lipidomics on the physiologically relevant aerobic-to-anaerobic transition in the presence of nitrate, a window in which respiratory modules are rapidly reprogrammed [24].

Quantitative LC-MS lipidomics first established how each synthase contributes to CL levels under these conditions (Figure S2). Strains carrying a Δ*clsA* mutation exhibited a marked reduction in CL abundance, highlighting ClsA as the dominant enzyme sustaining baseline CL content. In contrast, Δ*clsB* and Δ*clsC* strains retained CL levels comparable to WT, indicating a more limited contribution of these synthases in our growth regime (Figure S2). Most strikingly, the Δ*clsABC* mutant displayed a complete loss of detectable CL consistent with observations made under aerobic conditions (Figure S2) [17].

We then profiled glycerophospholipids and neutral-lipid species in WT and Δ*clsABC* cells and identified differentially abundant lipids relative to WT (Fig. 4A). CL depletion triggered extensive lipidome remodelling, with numerous species significantly altered (Fig. 4A). At the lipid-class level, Δ*clsABC* cells showed increased phosphatidylglycerol (PG) and phosphatidic acid (PA), accompanied by elevated diacylglycerol (DAG) (Fig. 4B). The increase in PG was expected and is consistent with prior work [38], as PG is the direct precursor used by ClsA to generate CL through condensation of two PG molecules. Loss of ClsA-dependent flux would therefore be predicted to divert lipid pools toward PG accumulation.

**Figure 4.**
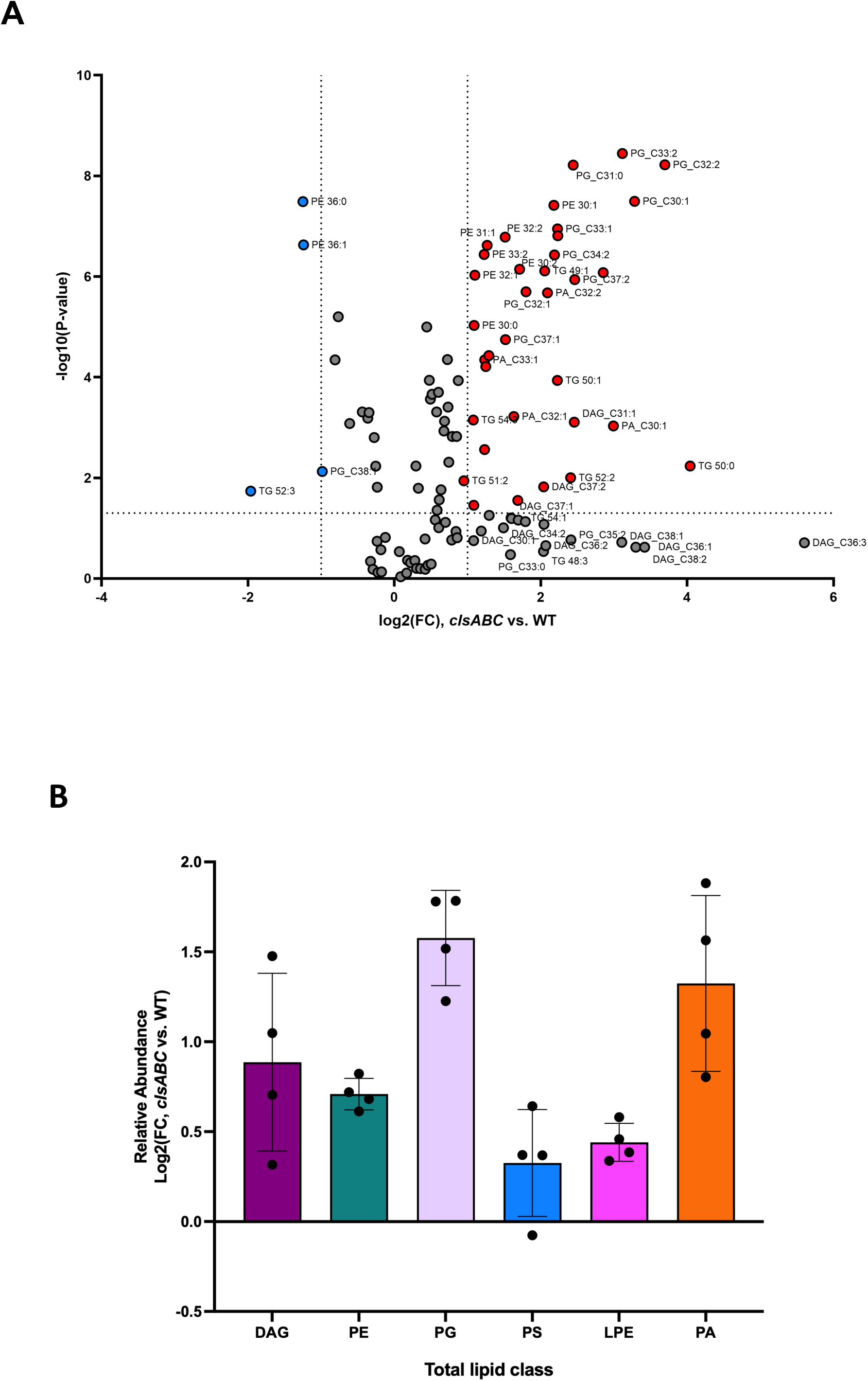
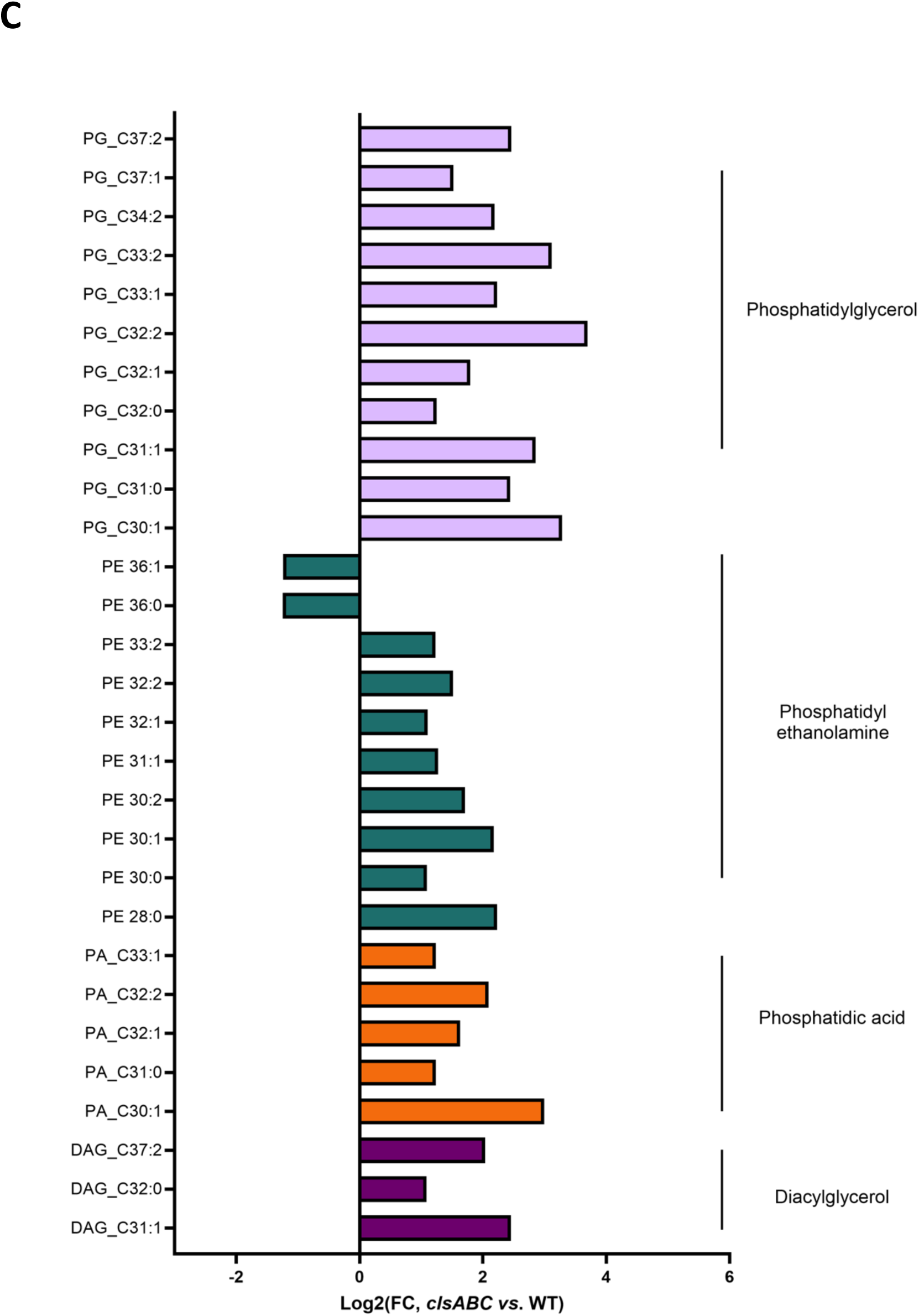
Differential lipid abundance between WT and Δ*clsABC* strains during transition from aerobic to anaerobic conditions. **(A)** Volcano plot showing the differential abundance of lipid species between WT and Δ*clsABC* strains. Lipids significantly enriched in the Δ*clsABC* mutant are highlighted on the right (red), while those enriched in WT are highlighted on the left (blue) (*P* < 0.05, *t*-test). Lipid species are represented as log₂ fold changes (log₂(FC)) versus - log₁₀(*P*-value), with data generated from four biological replicates and identified by LC-MS analysis. **(B)** Lipid classes are expressed as log₂ fold changes (log₂(FC)) of Δ*clsABC* versus WT and include: Diacylglycerol (DAG), Phosphatidylethanolamine (PE), Phosphatidylglycerol (PG), Phosphatidylserine (PS), Lysophosphatidylethanolamine (LPE), and Phosphatidic Acid (PA). Data are presented as mean ± SD. **(C)** Comparison of selected lipid species significantly altered (*P* < 0.05, *t*-test) between WT and Δ*clsABC* strains. Lipid species are categorized by class (TG, PG, PE, PA, and DAG) and displayed as bar plots of log₂ fold changes (log₂(FC)) for Δ*clsABC* versus WT.

Beyond these class-specific shifts, a salient and previously underappreciated trend emerged: CL depletion was associated with a broad increase in unsaturated fatty-acid (UFA) content across lipid classes, rather than being confined to a single lipid family (Fig. 4A and 4C). Collectively, these data show that CL depletion drives lipid rebalancing across the lipidome rather than an isolated loss of a single lipid class, and reveals a pronounced shift toward higher acyl-chain unsaturation, consistent with compensatory remodelling expected to increase membrane fluidity during nitrate-respiring condition.

### Cardiolipin depletion triggers system-wide proteome reprogramming during the nitrate-respiring transition

To understand how the nitrate-specific growth defect arises, we next investigated whether CL synthase mutations trigger broader proteome remodelling under nitrate-respiring conditions. Given that lipidomics revealed extensive membrane rewiring upon CL depletion, we hypothesized that loss of CL may engage stress responses and metabolic adaptations that reshape respiratory capacity and cellular homeostasis. To test this, we performed high-resolution MS-based quantitative proteomics on WT, each single mutant (Δ*clsA*, Δ*clsB*, Δ*clsC*) and the Δ*clsABC* strain during aerobic-to-anaerobic transition in the presence of nitrate.

Proteome profiles segregated according to the contribution of each synthase to CL production. Consistent with the lipidomics, Δ*clsB* and Δ*clsC* cells were essentially indistinguishable from WT, showing no detectable global proteome shifts (Fig. S4B and C). In contrast, Δ*clsA* and Δ*clsABC* displayed highly similar remodelling patterns, indicating that loss of ClsA-driven CL synthesis is sufficient to elicit the proteomic response in the growth regime tested (Figure 5 and Fig. S4A).

**Figure 5.**
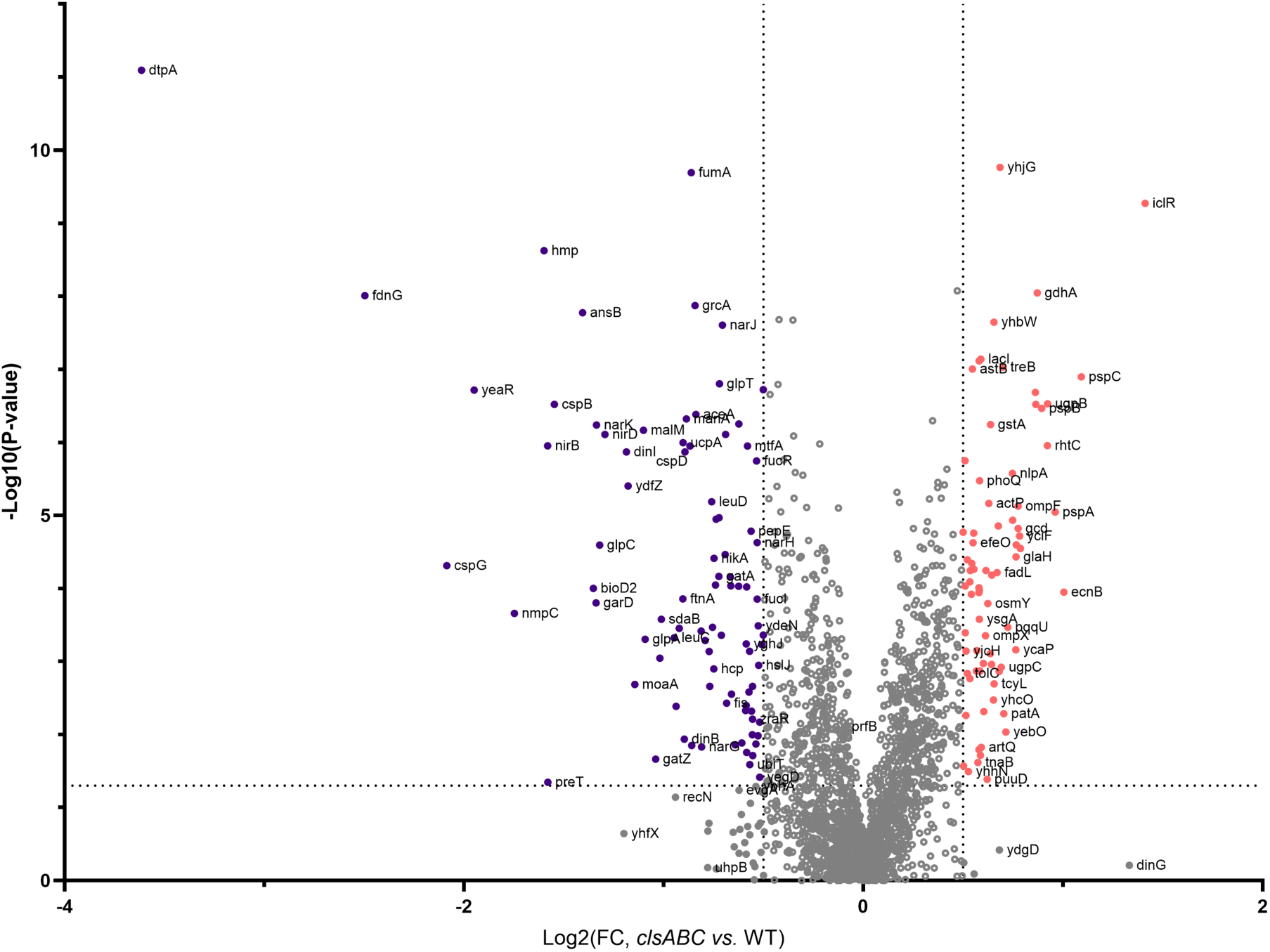
Cardiolipin depletion triggers system-wide proteome remodeling, linking nitrate respiration to NO homeostasis and membrane stress. Volcano plot depicting the differential expression of proteins between Δ*clsABC* and wild-type (WT) strains. The x-axis represents the log₂ fold change (Log₂(FC)) in protein abundance, with negative values indicating downregulation and positive values indicating upregulation in Δ*clsABC* compared to WT. The y-axis represents the - log₁₀(p-value), highlighting the statistical significance of the observed changes. Proteins with significant differential expression are colored in purple (downregulated) and red (upregulated), while non-significant proteins are shown in grey.

In the Δ*clsA* and Δ*clsABC* strains, proteins linked to nitrate respiration and reactive nitrogen stress management were broadly reduced (Fig. 5, Fig. S4A). This included the cytoplasmic nitrite reductase subunits NirB/NirD, the nitrate/nitrite transporter NarK, the NO-handling proteins Hmp and, to a lesser extent, Hcp. NirBD and Hcp are key contributors to NO control during nitrate respiration [34], and their decreased abundance suggests that the NO-homeostasis arm, which normally balances respiratory flux, is weakened in CL-deficient backgrounds. Additional decreases were observed in cold-shock/stress proteins (*e.g.*, CspB and CspG) and central metabolic enzymes (*e.g.*, FumA and GlpABC), further supporting a system-level reprogramming rather than an isolated defect in a single pathway. Importantly, although some respiration-associated factors shifted at the proteome level, the preserved Fdn activity and BV-supported Nar activity, together with the stability of tagged Nar and Fdn subunits, argue that the cellular pool of functional NarGHI and FdnGHI complexes is not grossly reduced; instead, CL deficiency primarily impacts quinol-linked Nar function and is accompanied by remodelling of accessory pathways engaged during nitrate respiration. In line with this broader view, 55% of the proteins whose abundance changed are not membrane-associated, and consistent with the reported links between CL/UFA and the Sec translocon [39–41], Sec-exported substrates showed abundance changes in our dataset.

Conversely, Δ*clsA* and Δ*clsABC* strains displayed induction of inner-membrane stress and compensatory programs, including higher levels of the phage shock proteins (PspA/PspB/PspC), alongside increased abundance of multiple stress and metabolic adaptation factors such as IclR or GdhA (Figure 5, Fig. S4). Collectively, these data demonstrate that impaired ClsA-dependent CL synthesis triggers coordinated proteome remodeling during the aerobic-to-anaerobic transition under nitrate-respiring conditions. This remodeling couples inner-membrane stress responses and metabolic reallocation to the reduced abundance of key nitrate-respiration and NO-control modules.

### Anaerobiosis drives lipid species remodelling toward higher unsaturation

To contextualize the CL-dependent phenotypes observed during the aerobic-to-anaerobic transition within a broader physiological framework, we conducted a quantitative analysis of the wild-type lipidome under three defined conditions: aerobic growth (ox+), aerobic-to-anaerobic transition (switch), and fully anaerobic nitrate-respiring growth (ox−) (Figure 6).

**Figure 6.**
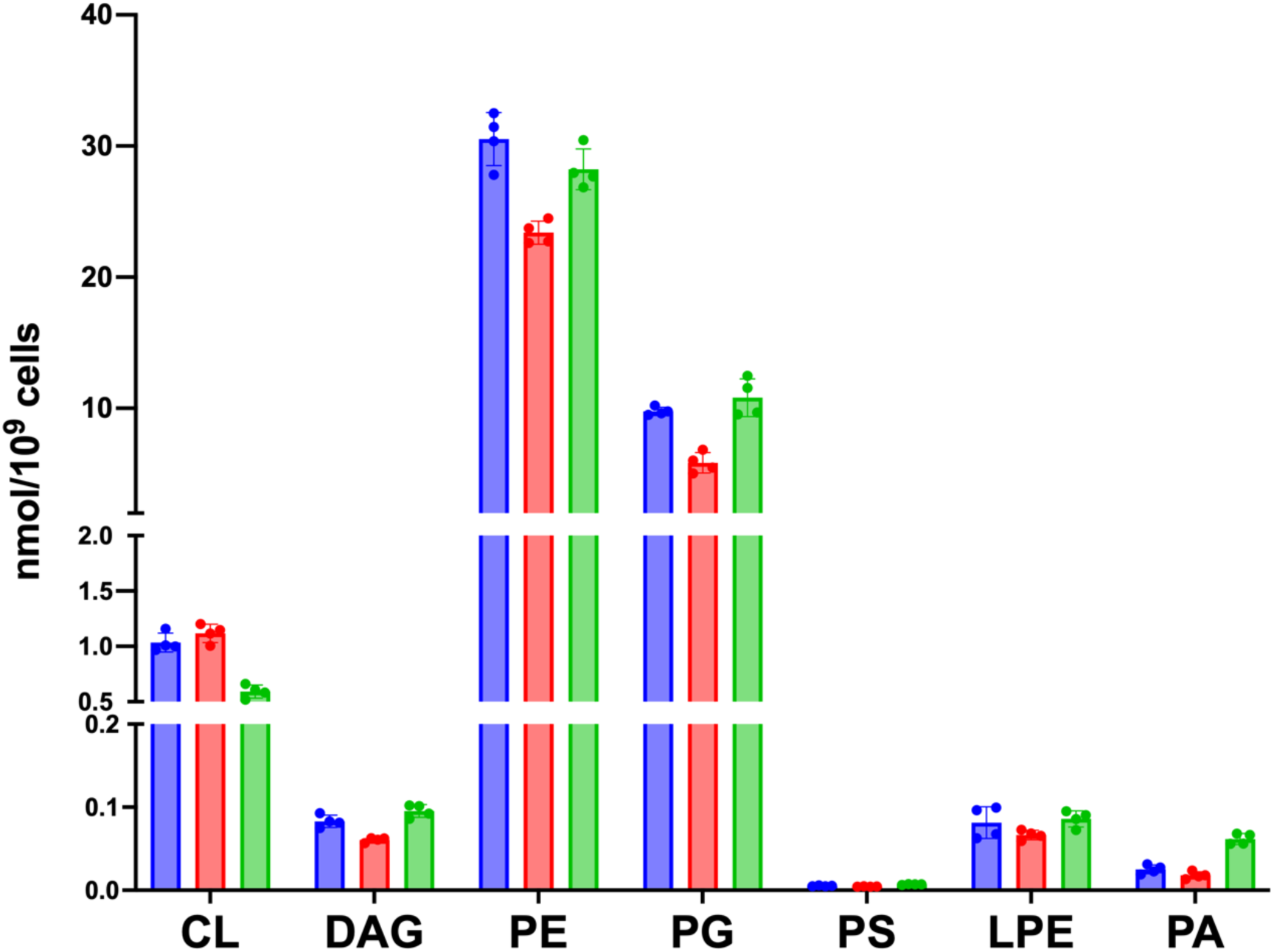
Remodeling of the *E. coli* wild-type strain lipidome during the aerobic-to-anaerobic transition and under established anaerobic conditions. Total lipid abundance levels expressed as nmol/10⁹ cells across three conditions: aerobic growth (ox+, blue), aerobic-to-anaerobic transition (switch, red), and nitrate-respiring anaerobic growth (ox−, green). Bars represent the mean lipid abundances measured from four biological replicates. Error bars indicate standard deviation (SD).

At the lipid class level, the transition state (switch) exhibited only minor changes relative to aerobiosis (Figure 6). The most notable class-level difference refers to a modest yet significant decrease in PE and PG in “switch” cells, while other major classes remained largely unchanged. In the anaerobic state (ox−), class-level remodelling remained subtle, with the most consistent change being a modest but significant decrease in CL. These observations suggest that the early stages of oxygen limitation do not drastically alter the lipid composition, but instead preserve overall class homeostasis throughout the transition. In contrast, species-level lipidomics separated the anaerobic endpoint (ox−) from both “ox+” and the transition state (switch) (data not shown). Volcano-plot patterns further indicated that the switch lipidome is more similar to “ox+” than to “ox−” (data not shown). The ox− shift was driven primarily by increased abundance of the most unsaturated lipid species. Together, these data show that, despite minimal class-level variation, lipid-species remodelling continues beyond the initial respiratory transition and consolidates only once nitrate-respiring anaerobic growth is established.

Together, these data identify a targeted membrane adaptation trajectory in wild-type cells. Early respiratory reprogramming involves minor class-level lipid changes, followed by extensive remodelling at the lipid molecular species level. This ultimately results in a distinct anaerobic lipid configuration characterized by a marked shift toward higher unsaturation.

## Discussion

Cardiolipin (CL) is a multifunctional phospholipid embedded in the bacterial membrane, where it influences a wide range of cellular processes, from respiratory efficiency to membrane homeostasis. While its role as a structural and functional modulator is well-documented, the precise mechanisms by which CL modulates membrane dynamics and cellular adaptation, particularly during metabolic transitions, remain incompletely understood. Here, we tackled this question by employing a multi-omics approach, integrating lipidomics, proteomics and genetic analysis, to investigate the consequences of CL depletion in *E. coli* under nitrate-respiring conditions. Our findings reveal that, under these conditions, CL depletion triggers a marked global rewiring of the *E. coli* lipidome. The most prominent signature is a strong enrichment of unsaturated fatty acids (UFAs) across lipid classes, together with increased phosphatidic acid (PA) and diacylglycerol (DAG), without a concomitant increase in the abundance of fatty-acid biosynthesis enzymes (Fig. S2). Similarly, the abundance of PlsB/PlsC is unchanged, indicating that PA/DAG accumulation is unlikely to result from increased canonical PA synthesis capacity, but instead from redistribution of lipid intermediates. Collectively, the observed system-level lipid remodelling reinforces the idea that CL biosynthesis is intricately linked to global lipid homeostasis rather than functioning as a simple terminal branch of lipid metabolism [42].

Our wild-type lipidome time course further refines this interpretation. At the transition state from aerobic to anaerobic growth, wild-type cells display only minor class-level changes relative to aerobiosis. In stark contrast, the Δ*clsABC* mutant already displays extensive lipidome rewiring at this same switch stage, characterized by a strong prevalence of more unsaturated lipid species. Strikingly, this tendency mirrors the physiological trajectory observed in wild-type cells: once anaerobic nitrate respiration is fully established, membranes also remodel primarily at the molecular-species level toward higher unsaturation, while overall class composition remains largely stable. Thus, CL depletion does not induce an arbitrary lipid state; rather, it amplifies a membrane remodelling trajectory that is part of the normal anaerobic adaptation program. This progressive, condition-dependent lipid remodeling aligns with observations in other bacteria, such as *Bacillus subtilis*, where hypoxic stress revealed that CL levels rapidly increase at the expense of its precursor PG, with CL accumulation correlating with enhanced cyanide-resistant respiration and altered membrane morphology [43]. This inverse relationship where *B. subtilis* upregulates CL in response to oxygen limitation while *E. coli* remodels its lipidome toward UFAs highlights species-specific strategies for maintaining membrane function under metabolic stress.

The directionality of these lipid changes is particularly informative. The observed increased of UFA content is expected to reduce acyl chain packing and increase bilayer fluidity, consistent with homeoviscous adaptation [44]. CL itself, however, is also linked to membrane properties and mesoscale organization, including curvature stress, elasticity, and lateral heterogeneity [45]. We therefore interpret the UFA enrichment as a compensatory response aimed at preserving membrane physical properties in the absence of CL. This interpretation aligns with recent observations in the gut anaerobe *Bacteroides fragilis*, where loss of CL biosynthesis also triggers acyl chain remodelling, including increased unsaturation, predicted to alter membrane fluidity and permeability [46]. This cross-species convergence supports the idea that CL depletion elicits a conserved membrane-level adaptation program, even if the precise lipid classes and acyl chain trajectories differ between lineages.

Despite this apparent membrane-level compensation, CL deficiency does not restore efficient nitrate-dependent energy conservation. Our data localize the functional defect to quinol-linked catalysis rather than to complex abundance, biogenesis, or intrinsic activity of the catalytic module. This pattern is consistent with CL affecting the membrane-embedded quinol oxidation interface of NarGHI. Structural evidence supports such a mechanism: CL has been observed at defined sites in NarI, with an unsaturated acyl chain in close proximity to heme *b*_D_ and one its ligand altering the EPR line shape of heme *b*_D_, suggesting structural modulation at the heme center [21]. In this framework, CL does not merely provide generic negative charge; instead, it acts as a privileged lipid cofactor that stabilizes a productive geometry for quinol engagement and electron transfer through the hemes. While the increase in UFAs may enhance membrane plasticity, it fails to fully recapitulate the specific CL-defined microenvironment required for optimal quinol-linked turnover and coupling efficiency. This underscores a critical distinction between restoring bulk membrane fluidity and reconstructing the precise lipid-protein interaction landscape necessary for respiratory function.

Our proteomics analysis further positions CL at the intersection of nitrate respiration and nitrosative-stress control. CL deficiency is associated with reduced NarK abundance and lower levels of NO-homeostasis proteins (NirBD, Hmp, and Hcp). This pattern is consistent with two non-exclusive scenarios: (*i*) reduced nitrate import and impaired quinol-linked Nar activity decrease nitrite/NO production, thereby lowering demand for NO-detoxification systems; or (*ii*) altered respiratory flux and electron partitioning increase local nitrosative stress, but this is not resolved by steady-state protein abundances alone. Importantly, under our conditions, CL deficiency does not exacerbate the pronounced cell-length heterogeneity intrinsically associated with nitrate respiration, arguing against a major increase in NO-associated morphological stress.

In parallel, CL loss correlates with increased abundance of membrane stress proteins, including Psp components, consistent with compensatory mechanisms that preserve inner-membrane function during lipidome remodelling. Notably, we observe no clear signatures of generalized envelope or cytoplasmic stress, contrasting with aerobic studies that report growth delays and stress pathway activation in CL-deficient cells under different media conditions [42]. This suggests that the metabolic state and nutrient environment critically shape the cellular consequences of CL depletion.

Consistent with this context dependence, we detect no major morphology phenotype upon CL loss under our anaerobic conditions. CL-driven morphology changes have been reported in a species- and condition-dependent manner, including shortened cells in *Rhodobacter sphaeroides* [47] and altered dimensions in other lineages, whereas early studies in *E. coli* and *B. subtilis* found no comparable shape alterations [48,49]. More recent work revisiting *E. coli* reported enlarged periplasmic space at cell poles and increased length/width under specific aerobic rich-media conditions [42,50]. Our findings support the emerging view that morphological phenotypes are not universal consequences of CL loss but may arise when CL depletion intersects with specific envelope stresses or growth regimes, conditions not strongly engaged in our experiments.

Finally, our results are best interpreted within ecologically realistic oxygen landscapes. In its natural niches, *E. coli* experiences spatial and temporal oxygen gradients and repeatedly transitions between aerobic, microaerobic, and anaerobic states, often with fluctuating nitrate availability [25]. The aerobic-to-anaerobic transition analysed here therefore reflects an ecologically meaningful window where respiratory modules and their membrane environment co-remodel. Although complete loss of major phospholipids is not itself a typical physiological state, envelope lipid levels can vary dramatically in response to host-associated challenges [51], and bacterial surface remodelling is a hallmark of host-pathogen interactions [52]. In this broader context, our study identifies CL biosynthesis as a determinant of nitrate respiration efficiency and associated stress physiology during oxygen limitation, providing a mechanistic route from lipidome remodelling to altered respiratory performance *in vivo*.

## Material and methods

### Bacterial strains and growth conditions

The *E. coli* strains used in this study are derivates of RK4353 and listed in the supplementary material (Table S1). *E. coli* strains were grown in LB medium or on LB agar plates for genetic construction, transformation, and storage. For all the various assays, bacteria were grown aerobically at 37°C in a chemically defined medium supplemented with 140 mM glycerol used as the main carbon source and 100 mM nitrate or fumarate as terminal electron acceptors. Anoxic growth of bacteria is performed in gas-tight Hungate tubes under Ar atmosphere or in a microplate reader (Spark Multimode Microplate Reader, TECAN®) placed within an anaerobic chamber. For anoxic fermentative growth, glycerol was replaced by 40 mM glucose and nitrate was omitted. To initiate the aerobic-to-anaerobic transition, *E. coli* cultures were first grown aerobically in glycerol-nitrate medium to an OD_600_ of 0.6, then transferred to gas-tight Hungate tubes. Immediately after transfer, the medium was sparged with argon to remove dissolved oxygen, and the tubes were sealed under an Ar atmosphere to establish anaerobic conditions. Cultures were then incubated anaerobically for 1.5 hours prior to further analysis or experimentation. The chemically defined medium is composed of potassium phosphate buffer (100 mM; pH 7.4), ammonium sulfate (6 mM), NaCl (9 mM), magnesium sulfate (2 mM), sodium molybdate (5 µM), sodium selenite (1 µM), Mohr’s salt (10 µM), calcium chloride (100 µM), Casamino Acids (0.5%), and thiamine (0.01%). Antibiotic were added when required.

### Molecular technics

Molecular biology experiments were performed according to standard procedures and the suppliers recommendations. NucleoSpin Plasmid, Mini Kit (Macherey-Nagel) and GeneJET Genomic DNA purification Kit (ThermoFisher) were used for plasmid preparations and *E. coli* genomic DNA extractions, respectively. PCR was performed with either GoTaq DNA polymerase (Promega®) or PrimeSTAR Max DNA polymerase (Takara) when the product required high-fidelity polymerase.

### Construction of *E. coli cls* mutants and NarG-Halo_Tag_ and FdnI-Snap_Tag_ fusion strains

P1 transduction was used to transfer the *clsA::kn*, *clsB::kn*, *clsC::kn*, and *fdhE::kn* mutations from corresponding Keio collection strains into the JCB4023 strain. The transductants were purified twice on LB plates supplemented with kanamycin (50 µg/ml). The kanamycin cartridge was eliminated using pCP20 plasmid. Mutant genotypes were verified by PCR amplification using primers flanking the *clsA*, *clsB*, *clsC*, *fdhE* genes [*clsA*(fwd)/*clsA*(rev), *clsB*(fwd)/*clsB*(rev), *clsC*(fwd)/*clsC*(rev), and *fdhE*(fwd)/*fdhE*(rev)]. The nitrate-reductase-deficient strain JCB4023 was also used as recipient for integration of the translational *fdnI*-SNAP_Tag_ fusion at the natural chromosomal locus using λ-red mediated recombination method as described in [53] and complemented with the low-copy number plasmid pVA70-*narG::HT* resulting in production of Halo-tagged NarGHI under the control of its native promoter. SNAP_Tag_ was amplified from plasmid pUC::SNAP_Tag_ (Genecust) using primers *SNAP*(fwd) and *SNAP*(rev). The kanamycin resistance gene (*kn*) was amplified from genomic DNA purified from a Keio collection strain using primers *kn*(fwd) and *kn*(rev). The *fdnI-SNAP::kn* fragment used for the recombination was amplified from the above-described SNAP and the *kn* fragments using primers *fdnI-SNAP*(fwd) and *kn*(rev), which imparted flanking homologous regions to the chromosomal *fdnGHI* operon. The kanamycin-resistant recombinant LCB4720 was purified twice on LB plates supplemented with kanamycin (50 µg/ml) and characterized by PCR using primers *fdnI*(fwd) and *fdnI*(rev) and sequencing. The same strategy was employed for the construction of the *fdnI-Halo_Tag_* strain. The Halo_Tag_ was amplified from plasmid p*UC::Halo_Tag_* (Genecust) using primers *Halo*(fwd) and *Halo*(rev), and the *fdnI-Halo_Tag_:kn* fragment used for the recombination was amplified from the above-described SNAP and kn fragments using primers *fdnI-Halo_Tag_*(fwd) and *kn*(rev), which imparted flanking homologous regions to the chromosomal fdnGHI operon.To generate a plasmid enabling the production of a Halo-tagged NarGHI complex, the *Halo-tag* gene was amplified by PCR from the pUC::Halo_Tag_ vector (Genecust) using primers *Halo*(fwd) and *Halo*(rev), which introduced flanking regions homologous to the 3′ end of narG and the pVA70 plasmid backbone. In parallel, the pVA70 plasmid was linearized by PCR to create homologous overhangs compatible with the Halo_Tag_ fragment and narG flanking regions [*pVAflankA(*fwd)/*pVAflankB*(rev)]. The amplified *Halo-tag* fragment and linearized plasmid were assembled using Gibson Assembly (NEB), yielding the recombinant plasmid p*VA-narG::HT* and purified on LB plates supplemented with Amp (100 µg/ml). This construct enables the in-frame fusion of the *Halo-tag* to the C-terminus of *narG*, allowing for the production of a Halo-tagged NarGHI complex. Oligonucleotides used in this work are listed in Table S3.

### Mass spectrometry based-Lipidomic analyses on purified protein

*E. coli* cells were metabolically quenched in dry ice - ethanol for 1 min then washed three times in cold phosphate-buffered saline. Total lipid was extracted from a cell suspension or from two negative control proteins that do not bind lipids (namely, catalase 1 and catalase 2). The extraction process utilized 350 µL chloroform/methanol (4:3 v/v) containing 1 nmol each of internal standards (as listed in the following table, Avanti-Sigma), followed by the addition of 140 µL of water to induce biphasic separation. The lower organic phase was collected and subsequently dried in a vial insert. The recovered lipids were dissolved in methanol (50 µL) and aliquot (10 µL) was injected to a Agilent 1290 infinity/Infinity II LCMS system equipped with ZORBAX Eclipse Plus C18, 100 x 2.1 mm, 1.8 um reversed-phase column (Agilent) with Infinity II inline filter, 0.3 um (Agilent) maintained at 45C. Samples were separated by the change in gradient of solvent A (Water:acetonitrile:isopropanol, 5:3:2 (v/v/v), 10 mM ammonium formate) and solvent B (Isopropanol:acetonitrile:water, 90:9:1 (v/v/v), 10 mM ammonium formate). The gradient was as follows, starting with a flow rate of 0.4 mL/min at 15% B and increasing to 50% over 2.5 min then to 57% at 2.6 min, then to 70% over 9 min and to 93% at 9.1 min, then to 96% over 11 min and to 100% at 11.1 min and hold to 12 min then back to 15% at 12.2 min to 16 min (total of 16 min). MS (Agilent 6495c triple quadrupole) was operated in for targeted analysis with DMRM (dynamic multiple reaction monitoring) with 650 ms of cycle time with following setting: Agilent Jet stream ion source with positive and negative switching with gas temperature at 150°C, drying gas (nitrogen) 17 L/min, Nebulizer gas 20 psi, Sheath gas temperature at 200°C with flow at 10 L/min, capillary voltage 3500 V for positive and -3000 V for negative mode, nozzle voltage at 1000 V for positive and -1500 V for negative mode. As described [54].For targeted lipidomic analysis *E. coli*, first, collision energy (CE) and characteristic fragmentation for Lysil-phosphatidylglycerol (LyPG), was obtained by Mass Hunter Optimiser software (Agilent) using authentic standard LyPG C18:1 (Sigma). Then the fatty acid combination for each lipid species was confirmed by *E. coli* total lipid using neutral loss ion scan (NL m/z 35 for DAG with CE 35 V; m/z 141 for LPE and PE with CE 19 and 17 V, respectively; m/z 300 for LyPG with CE 41 V; m/z 189 for PG with CE 19 V, m/z 115 for PA with CE 13 V in positive mode). For CL, PG species detected as above was further used to determine CL species in *E. coli*. The resulting lipidomic quantitative data were further normalized using flow cytometry values to ensure accurate cross-sample comparability and LCMS data was combined to pre-existing dynamic multiple reaction monitoring (DMRM) method for optimized targeted lipidomic analysis for *E. coli*.

### Bacterial Enumeration Using Flow Cytometry

Bacteria were enumerated using flow cytometry. For each condition, 1 mL of bacterial culture was collected, pelleted, and fixed in 1 mL of PBS containing a final 0.25% glutaraldehyde, 0.01% pluronic solution. Fixed samples were stored at 4°C until analysis. For flow cytometry analysis, samples were thawed at room temperature, diluted 100-fold, nucleic acid were stained II (Molecular Probes, commercial solution diluted 10 times) with SYBR Green for 15 minutes in the dark [55]. Acquisition was performed using a CytoFLEX Legacy analyzer (Beckman Coulter) at the PRECYM flow cytometry platform (https://precym.mio.osupytheas.fr/). The analyzer is equipped with three solid-state lasers (488 nm, 405 nm, and 638 nm) and a peristaltic pump system for precise measurement of the absolute volume analyzed. Data were acquired in log scale and stored in list mode, and subsequent bacterial enumeration was analyzed *a posteriori* using CytExpert 2.6 software (Beckman Coulter).

### Measurement of enzyme activity

Bacterial cells grown under nitrate-respiring conditions were harvested at mid-exponential phase, washed, and resuspended in a buffer containing 40 mM Tris-HCl (pH 7.4) and 1 mM MgCl2. Bacterial cells were broken by one passage through a French press, followed by a gentle centrifugation at 5,000 rpm for 10 minutes to remove cell debris. The supernatant was then aliquoted, frozen in liquid nitrogen, and stored at -80°C until use.

Nitrate reductase activity was measured with standard assays using reduced benzyl viologen or menadiol as electron donors, in cells harvested during the aerobic-to-anaerobic transition. One unit of nitrate reductase activity is expressed in micromoles of nitrate reduced per minute per milligram of total proteins [31,56]. For formate dehydrogenase activity, cells were harvested from anaerobic growth and the activity was measured by 2,6-dichlorophenolindophenol (DCPIP) reduction mediated by phenazine methosulfate (PMS) as described previously [29]. One unit of formate dehydrogenase activity is expressed in micromoles of formate oxidized per minute per milligram of total proteins. Total protein content was quantified using the Bradford method, with Bovine Serum Albumin (ThermoFisher) employed as the calibration standard.

### Western blot assay

Western blots were performed with 100 µg of proteins loaded and separated on a 10% Mini-Protean TGX Stain-Free precast protein gel (BioRad) and subsequently electro-transferred on a polyvinylidene difluoride (PVDF) membrane (Biorad). The *E. coli* NarG-Halo_TAG_ and FdnI-SNAP_TAG_ proteins were detected with anti-NarG antibodies (dilution 1/2000, [32,34]) and anti-SNAP (dilution 1/500, CAB4255 ThermoFisher) respectively both produced in rabbit. HRP-anti-rabbit (A6154 Merck) was used as secondary antibodies at 1:2,000 dilution and the detection was achieved with the ECL reagent (ThermoFisher). A stain-free gel was used to precisely normalize the amounts of proteins in each lane of the gel used for the western.

### ATP assay

Bacterial strains were grown in a defined M9-glycerol medium supplemented with 10 mM sodium formate and 100 mM nitrate to mid-exponential phase and subsequently transferred to anaerobic conditions for 1.5 hours to induce dehydrogenase/reductase expressions. Cells were then harvested, washed, and resuspended in phosphate-based assay buffer (40 mM potassium phosphate, 10 mM magnesium sulfate, pH 7.0), followed by incubation at room temperature for 45 minutes in an anaerobic chamber to obtain a resting cell suspension. Reactions were initiated by mixing the cell suspension at a 1:1 ratio with assay buffer containing either sodium formate (10 mM), nitrate (100 mM), or both compounds in 1.5 mL tubes and were allowed to proceed for 10 minutes. At the end of the reaction period, 10 μL aliquots were withdrawn and transferred into a separate 96-well plate containing 90 μL of DMSO to terminate the reaction and release intracellular ATP. ATP levels were quantified using an ATP determination kit ([57], Invitrogen A22066). Luminescence was recorded using black 96-well plates (Corning 3915) on a microplate reader (Tecan Infinite M200) and calculated by comparison to a standard curve generated from known ATP concentrations.

### *In vivo* labelling of HaloTagged proteins and fluorescence microscopy

*E. coli* cultures expressing NarG or FdnI Halo_Tag_-fusion proteins were first grown aerobically in M9-glycerol medium supplemented with 140 mM glycerol and 100 mM nitrate at 37 °C to an OD₆₀₀ of 0.6. The same protocol for aerobic-to-anaerobic transition was applied, as previously described, to establish anaerobic conditions prior to further experimentation. After anaerobic incubation, 1 mL of culture was harvested by centrifugation at 5,000 × g for 2 min at room temperature (RT), and the pellet was resuspended in 100 µL of fresh M9-glycerol medium. HaloTag® TMR Ligand (Promega G8251) was added to a final concentration of 5 µM, and the mixture was incubated for 20 min at RT in an anaerobic chamber, protected from light [36,37]. Following labeling, cells were washed three times with 1 mL of M9-glycerol medium to remove unbound ligand, fixed with 4% formaldehyde in 1X PBS for 10 min at RT. All anaerobic steps were performed in an anaerobic chamber to maintain strict anoxic conditions. Two microliters of the suspension were mounted onto 1% agarose pads for fluorescence imaging as described in [34,35]. Imaging was performed using a Nikon Eclipse TiE PFS inverted epifluorescence microscope (100 oil objective, NA 1.3) and a Hamamatsu Orca Flash LT 4.0 sCMOS camera.

All image analysis and statistical representations of fluorescence were performed with FIJI and MicrobeJ softwares. The distribution of fluorescent clusters was obtained by a local maxima detection algorithm and reported to the relative longitudinal axis position; the signal was prefiltered by band-pass fast Fourier transform (FFT). The average distribution heat maps of Halo-tagged NarG and FdnI were obtained by the projection of the raw fluorescence of all cells in a group. Each cell shape and the associated fluorescence signal were previously morphed to the group mean shape using MicrobeJ v5.13k.

### Mass spectrometry, data analysis and quantitative proteomics processing

Protein extracts were loaded on NuPAGE™ 4–12% Bis–tris acrylamide gels according to the manufacturer’s instructions (Life Technologies) and run for 5 to 7 minutes (80V) until whole sample enter into top of the gel. Running of protein was stopped as soon as proteins stacked in a single band. Protein containing bands were stained with Imperial Blue (Pierce), cut from the gel and digested with high sequencing grade trypsin (Promega) before mass spectrometry analysis. Gel pieces were washed and distained using few steps of 100mM NH_4_HCO_3_. Distained gel pieces were shrunk with 100 mM ammonium bicarbonate in 50% acetonitrile and dried at RT. Protein spots were then rehydrated using 10mM DTT in 25 mM ammonium bicarbonate pH 8.0 for 45 min at 56°C. This solution was replaced by 55 mM iodoacetamide in 25 mM ammonium bicarbonate pH 8.0 and the gel pieces were incubated for 30 min at room temperature in the dark. They were then washed twice in 25 mM ammonium bicarbonate and finally shrunk by incubation for 5 min with 25 mM ammonium bicarbonate in 50% acetonitrile. The resulting alkylated gel pieces were dried at room temperature. The dried gel pieces were rehydrated by incubation in 25 mM ammonium bicarbonate pH 8.0 supplemented with 12.5 ng/µl trypsin (Promega) for 1h at 4°C and then incubated overnight at 37°C. Peptides were harvested by collecting the initial digestion solution and carrying out two extractions; first in 5% formic acid and then in 5% formic acid in 60% acetonitrile. Pooled extracts were dried down in a centrifugal vacuum system. Samples were reconstituted in 0.1% TFA 4% acetonitrile before mass spectrometry using an Orbitrap Fusion Lumos Tribrid Mass Spectrometer (ThermoFisher Scientific, San Jose, CA) online with a neoVanquish chromatography system (ThermoFisher Scientific). Peptides were separated on EasySpray column (50cm, ES75500) at 40°C using a two steps linear gradient (4–20% acetonitrile/H2O; 0.1% formic acid for 110 min and 20-32% acetonitrile/H2O; 0.1% formic acid for 10 min). An EASY-Spray nanosource was used for peptide ionization (1,900 V, 275°C). MS was conducted using a data-independent acquisition mode (DIA). Full MS scans were acquired in the range of m/z 375–1,500 at a resolution of 120,000 at m/z 200 and the automatic gain control (AGC) was set at 4.0 × 10E5 with a 50 ms maximum injection time. MS2 spectra were acquired in the Orbitrap with a resolution of 30,000, in the mass range of 200-1800 m/z after isolation of parent ion in the quadrupole and fragmentation in the HCD cell under collision Energy of 30%. DIA parent ion range was from 400 to 1000 m/z divided into 40 windows 16 Da wide and from 1000 to 1500 m/z divided into 10 windows 50 Da wide.

For protein identification and quantification, relative intensity-based label-free quantification (LFQ) was processed using the DIA-NN 1.8 algorithm [58]. Raw files were searched against the *E. coli* (strain K12) database (referent proteome UP000000625) extracted from UniProt (date 2025-04-03; 4402 entries) and implemented with a contaminant database [59]. The following parameters were used for searches: (*i*) trypsin allowing cleavage before proline; (*ii*) one missed cleavage was allowed; (*iii*) cysteine carbamidomethylation (+57.02146) as a fixed modification and methionine oxidation (+15.99491) and N-terminal acetylation (+42.0106) as variable modifications; (iv) a maximum of 1 variable modification per peptide allowed; and (v) minimum peptide length was 7 amino acids and a maximum of 30 amino acids. The match between runs option was enabled. The precursor false discovery was set to 1%. DIA-NN parameters were set on double-pass mode for Neural Network classifier, Robust LC High precision for quantification strategy and RT-dependent mode for Cross-run normalization. Library was generated using Smart profiling set up. MS1 and MS2 mass accuracy was automatically calculated for precursor charge fixed between 2 and 4. The resulting label-free quantification (LFQ) data were further normalized using flow cytometry values to ensure accurate cross-sample comparability.

Main output file from DIA-NN was further filtered at 1% FDR and LFQ intensity was calculated using our DIAgui package at 1% q-value (https://github.com/marseille-proteomique/DIAgui [60]). The statistical analysis was done with Perseus software (version 1.6.15.0), where proteins were pre-filtered to remove contaminants ([61]. Relative quantification was calculated using fold changes of LFQ intensity between different conditions. To determine whether a given detected protein was specifically differential, a two-sample t-test was done using permutation-based FDR-controlled employing 250 permutations. The mass spectrometry proteomics data have been deposited to the ProteomeXchange Consortium via the PRIDE partner repository.

### Statistical analysis

The Student t-test was used to determine significant differences of the means of the data for all the experiments. Error bars represent the standard deviation, with * (P < 0.05), ** (P < 0.01), *** (P < 0.001), **** (P < 0.0001) indicating that the mean values are significantly different and ns that they are not significantly different (P > 0.05).

## Acknowledgements

We thank the Regional Flow Cytometry Platform for Microbiology (PRECYM) of the Mediterranean Institute of Oceanography (MIO) for providing the technical means and skills of its personnel, making available equipment and protocols, and methodological advice. PRECYM was supported by the European Regional Development Fund (ERDF; project 1166-39417) and by the French Groupement d’Intérêt Scientifique “Infrastructures en Biologie, Santé et Agronomie” (GIS IBiSA), which supports national research infrastructures. The GEMELI lipidomics platform was supported by the French National Research Agency (ANR-21-CE44-0010, ANR-23-CE15-0009-01, ANR-24-CE15-2171-02, ANR-25-CE30), the Fondation pour la Recherche Médicale (FRM EQU202103012700), Laboratoire d’Excellence Parafrap, France (ANR-11-LABX-0024), LIA-IRP CNRS Program (Apicolipid project), the Université Grenoble Alpes (IDEX ISP Apicolipid), Région Auvergne Rhone-Alpes (Grant IRICE Project GEMELI), labeled by IBiSA and the Collaborative Research Program Grant CEFIPRA (MESRI-DBT, Project 6003-1). Marseille Proteomics (marseille-proteomique.univ-amu.fr) was supported by IBISA (Infrastructures Biologie Santé et Agronomie), Plateforme Technologique Aix-Marseille, the Cancéropôle PACA, the Provence-Alpes-Côte d’Azur Region, the Institut Paoli-Calmettes, Fonds Européen de Développement Regional (FEDER) and Plan Cancer. This work was also supported by the French National Research Agency (ANR-21-CE44-0026-01-AM), the CNRS and Aix-Marseille University. M. Denic was funded by the ANR D2OX (ANR-21-CE44-0026-01-AM) project, M. Delprat was funded by the ANR ANAEROP (ANR-23-CE15-0017-02-AM) project. This project has received funding from Excellence Initiative of Aix-Marseille University - A*MIDEX, a French “Investissements d’Avenir” programme (AMX-19-IET-002).The funders had no role in study design, data collection and analysis, decision to publish or preparation of the manuscript.

---

**Figure S1.**
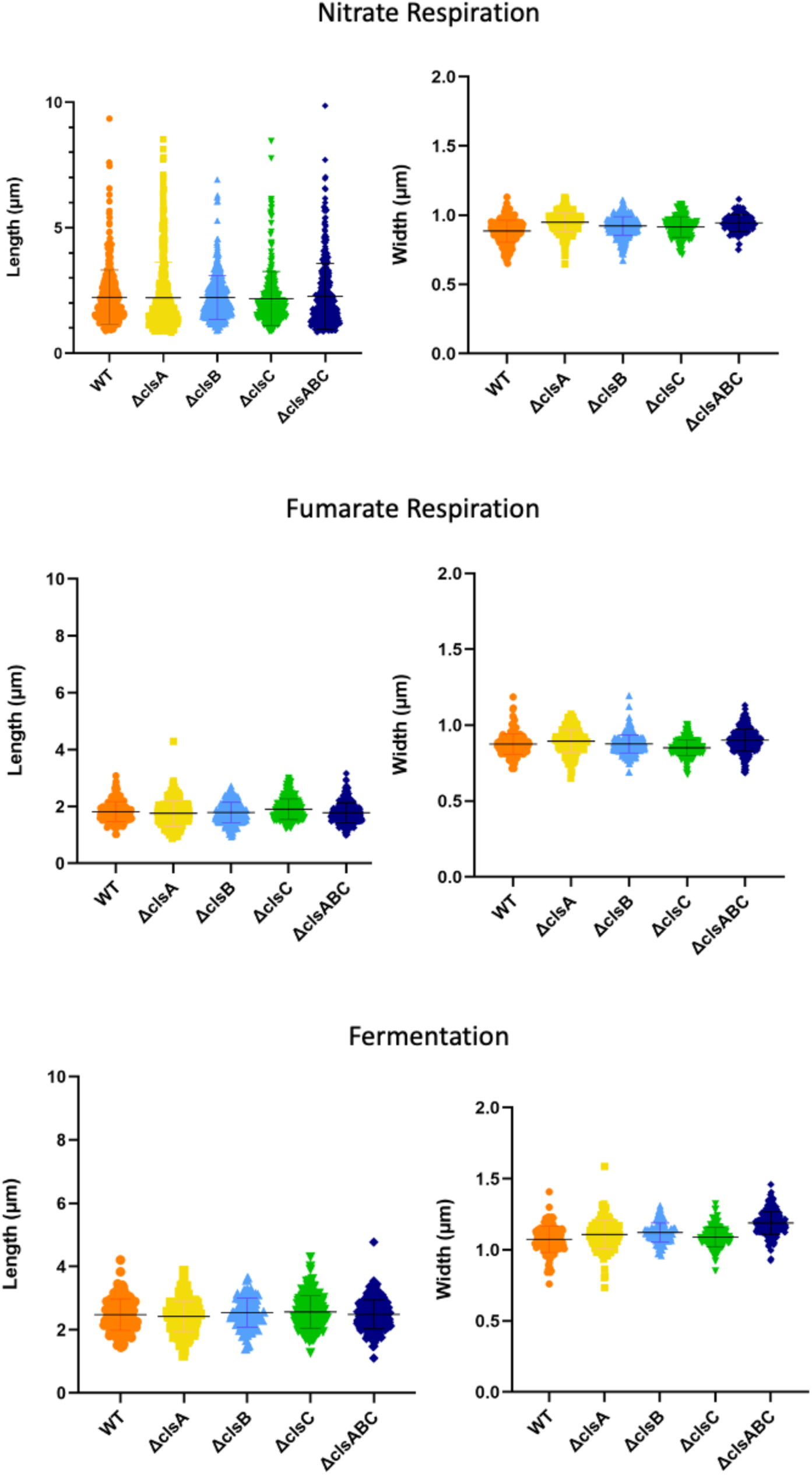
Growth and morphological analysis of wild-type and cardiolipin synthase mutant strains under different respiratory conditions. Violin plots depicting cell length and width measurements of WT, *clsA*, *clsB*, *clsC*, and *clsABC* mutant strains during nitrate respiration, fumarate respiration, and fermentation. Each plot shows the distribution of cell dimensions, with individual data points overlaid, n=300 cells.

**Figure S2.**
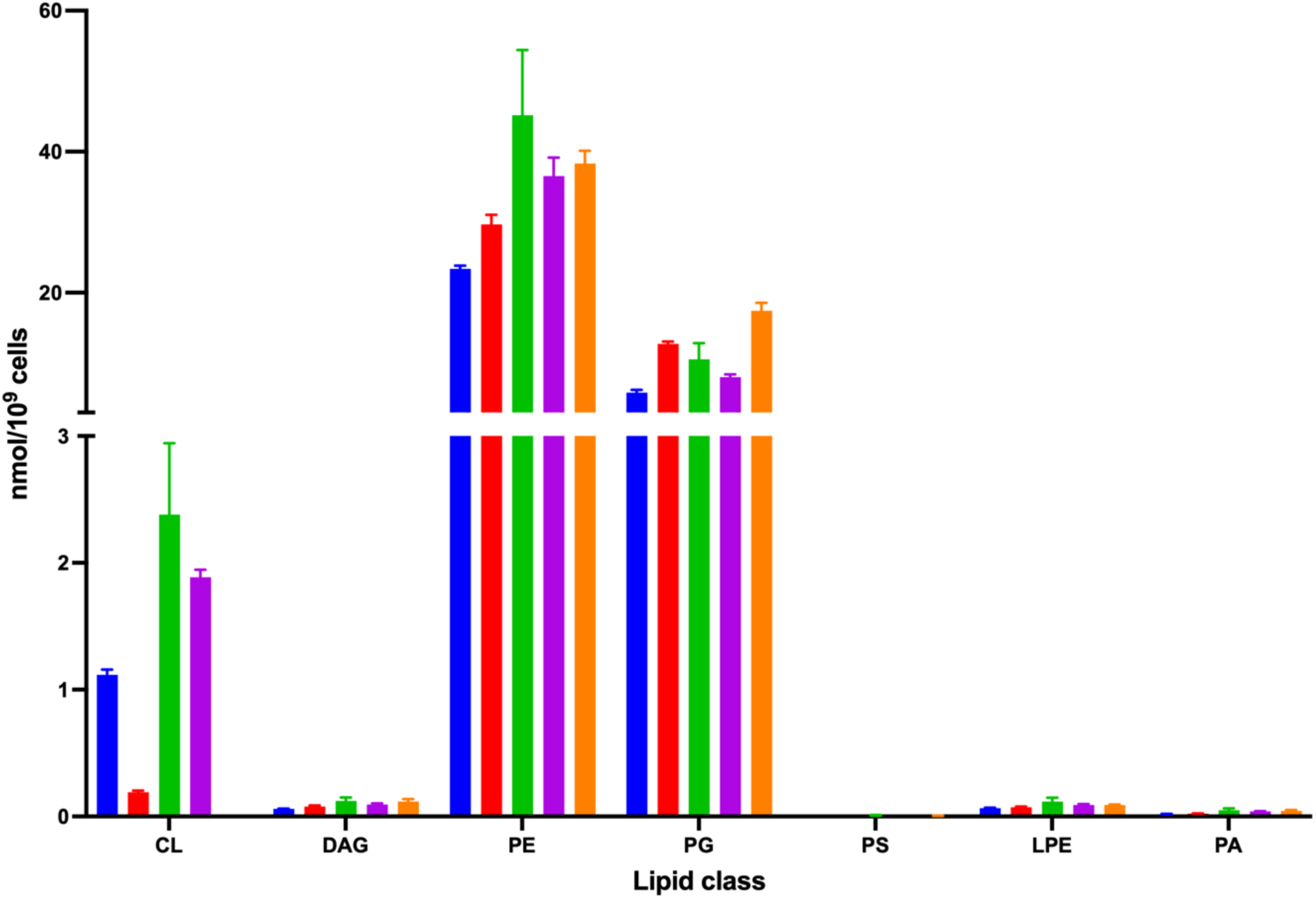
Lipidomic profiling reveals alterations in Δ*clsA* and Δ*clsABC* mutants compared to WT strains in aerobic-to-anaerobic transition. Bar graph illustrating the absolute abundance of major lipid classes in wild-type (blue) and cardiolipin synthase mutant strains Δ*clsA* (red), Δ*clsB* (green), Δ*clsC* (purple), and Δ*clsABC* (orange). The y-axis represents the lipid quantity (nmol per 10⁹ cells) for each lipid class: cardiolipin (**CL**), diacylglycerol (**DAG**), phosphatidylethanolamine (**PE**), phosphatidylglycerol (**PG**), phosphatidylserine (**PS**), lysophosphatidylethanolamine (**LPE**), and phosphatidic acid (**PA**). Error bars indicate standard deviation.

**Figure S3:**
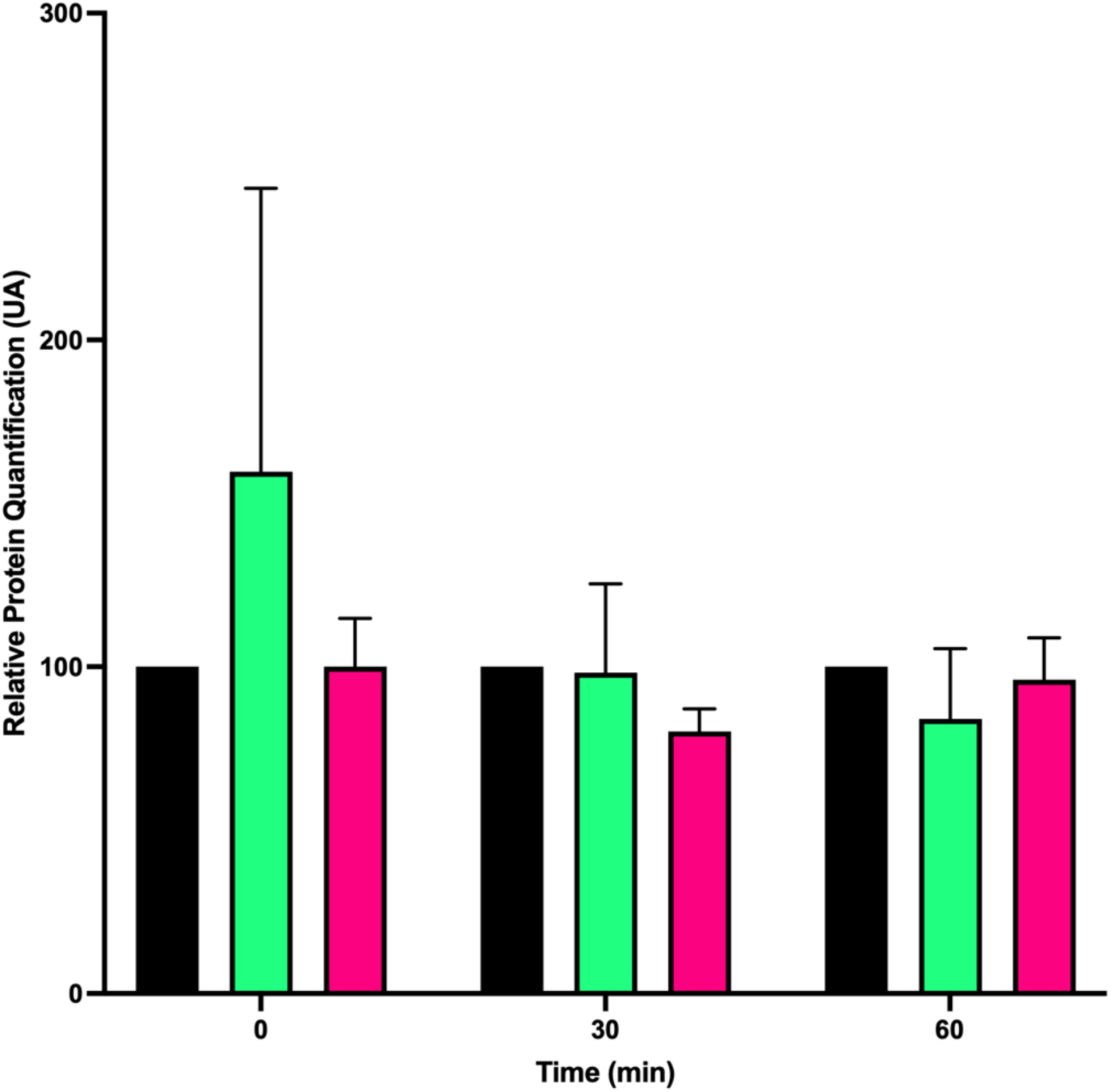
Quantification of protein stability during a chloramphenicol chase assay in WT and *clsABC* mutant strains. Quantification of protein stability during a chloramphenicol chase assay conducted at a concentration of 10 µg/ml. The black bars represent 100% of FdnI or NarG in the WT strain, serving as the baseline reference at each time point. The green and magenta bars show the relative abundance of FdnI and NarG, respectively, in the *clsABC* mutant background, enabling direct comparison to WT. Protein levels were monitored at 0-, 30-, and 60-minutes post-chloramphenicol treatment, with data representing the average of three biological replicates (n=3). Error bars indicate the standard deviation, illustrating the variability among replicates.

**Figure S4.**
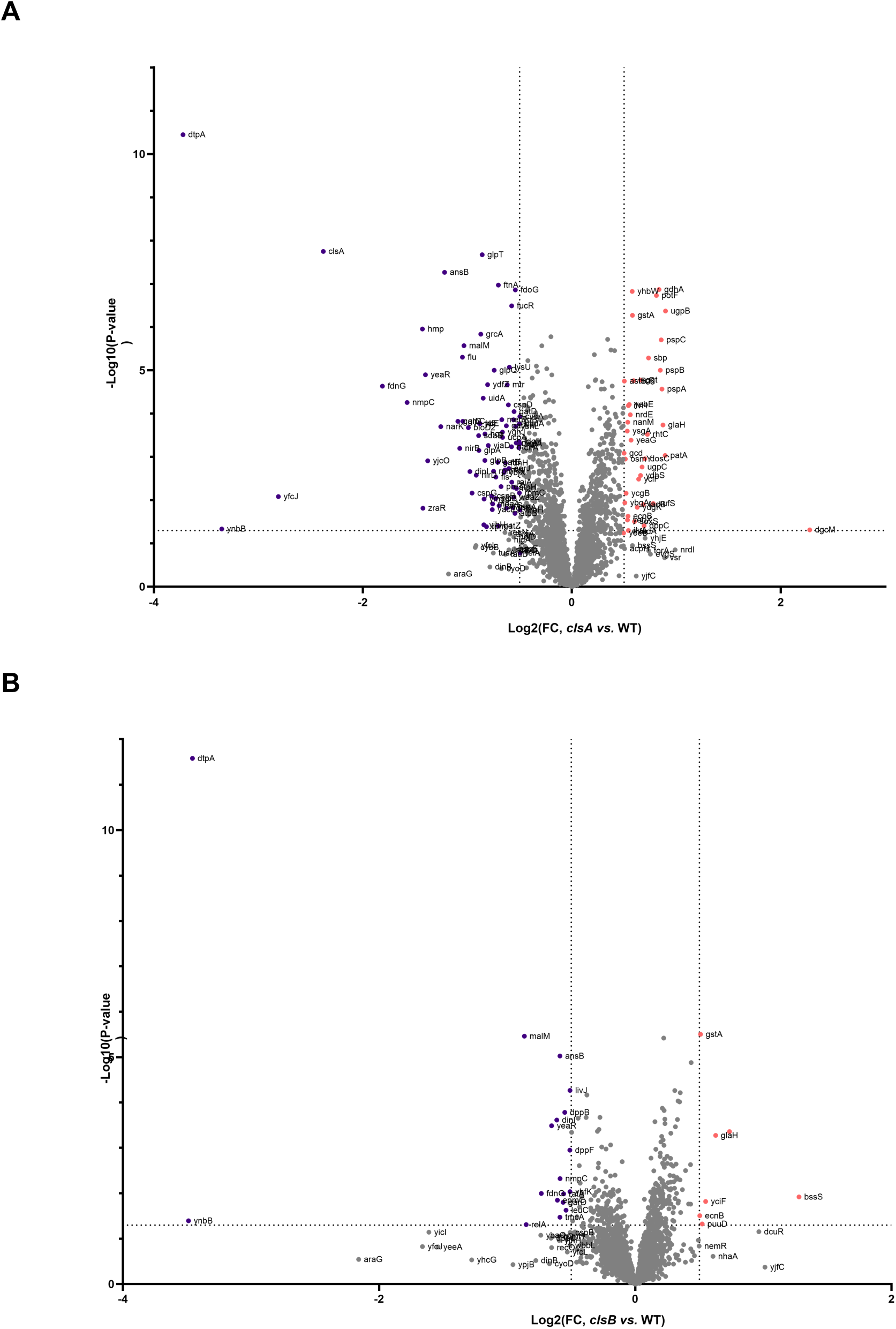
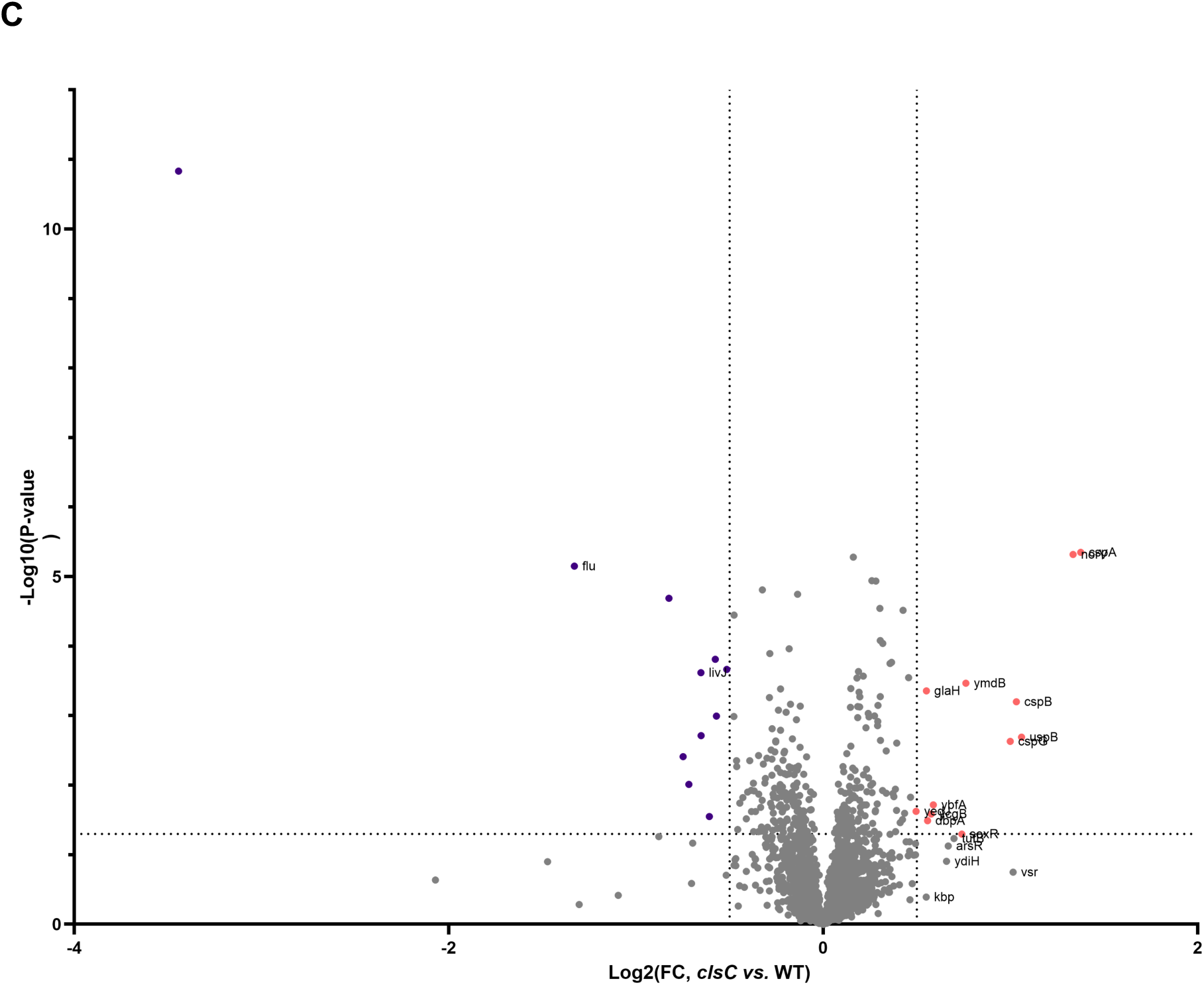
Comparative proteomic analysis of cardiolipin synthase individual mutants vs. WT strains. Volcano plot illustrating the differential protein abundance between *clsA* **(A)**, *clsB* **(B)** and *clsC* **(C)** mutants and WT strains, respectively. The x-axis represents the log₂ fold change (Log₂(FC)) in protein abundance, with negative values indicating downregulation and positive values indicating upregulation in the *clsA*, *clsB* or *clsC* mutant. The y-axis represents the -log₁₀(p-value), indicating the statistical significance of the changes. Proteins that are significantly altered are labeled.

**Table S1:**
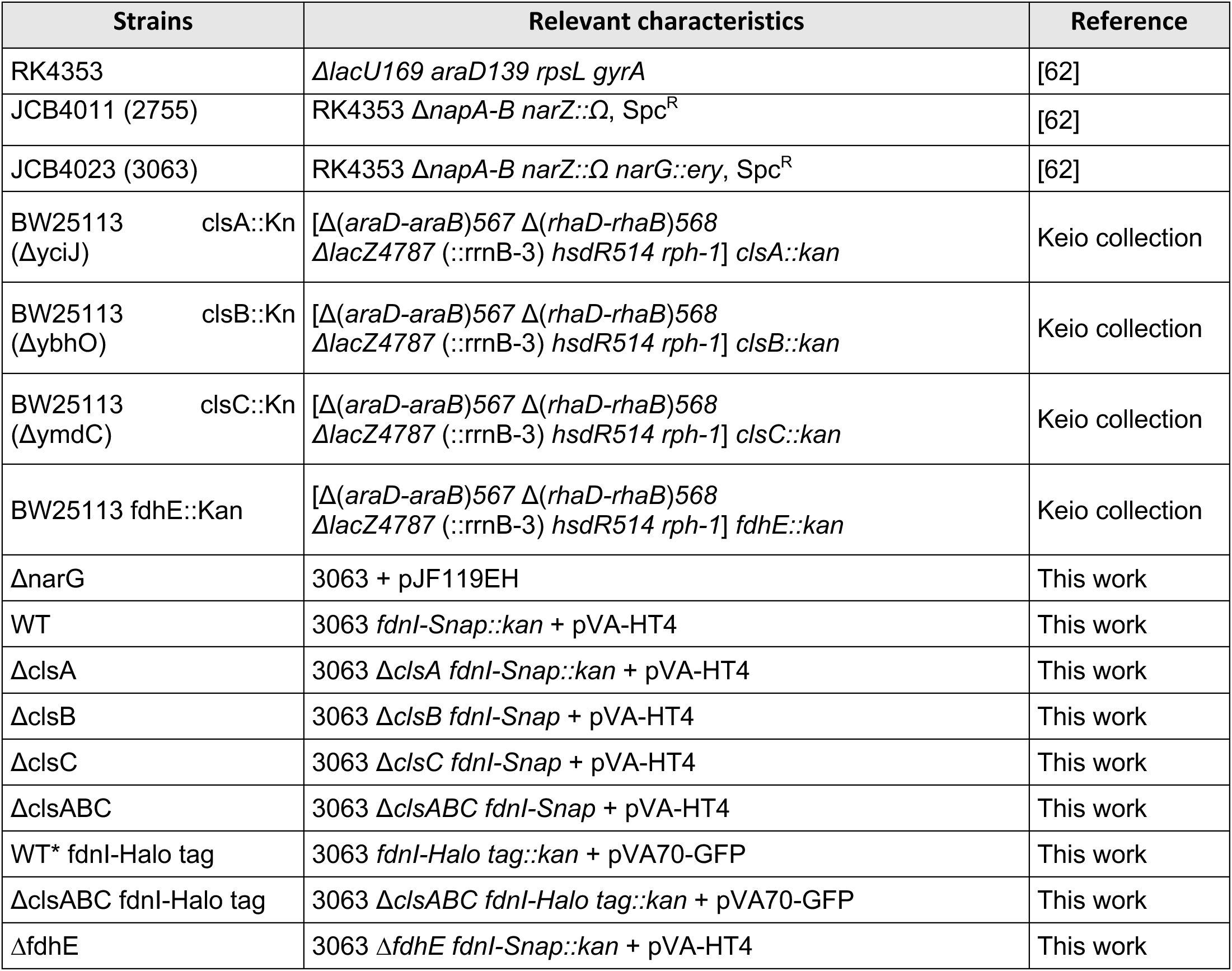
Bacterial strains used in this study.

**Table S2:** Plasmids used in this study.

| Plasmids | Relevant characteristics | Reference |
| --- | --- | --- |
| pUC::Snap tag | SNAP tag optimized codon | Genecust® |
| pUC::Halo tag | Halo tag optimized codon | Genecust® |
| pCP20 | *FLP*<sup>+</sup>, *λ*cl857<sup>+</sup>, *λP*<sub>R</sub> Rep<sup>ts</sup>, Ap<sup>R</sup>, Cm<sup>R</sup> | [53,63] |
| pJF119EH | *tacP*, *rrnB*, $\Delta$[*Xma*III-*Bam*HI, 0.5 kb], $\Omega$[*lacI*<sup>Q</sup>, *Hind*III, 1.2 kb]<br>$\Delta$[polylinker M13mp8], $\Omega$[polylinker M13mp18], Ap<sup>R</sup> | [35] |
| pVA-HT4 | pJF119EH, *P*<sub>nar</sub>-(*narG-Halo tag,narHJI*), Ap<sup>R</sup> | This work |
| pVA70 | pJF119EH, *P*<sub>nar</sub>-(*narGHJI*), Ap<sup>R</sup> | [35] |
| pVA70-GFP | pJF119EH, *P*<sub>nar</sub>-(*narG-egfp,narHJI*), Ap<sup>R</sup> | [35] |

**Table S3:**
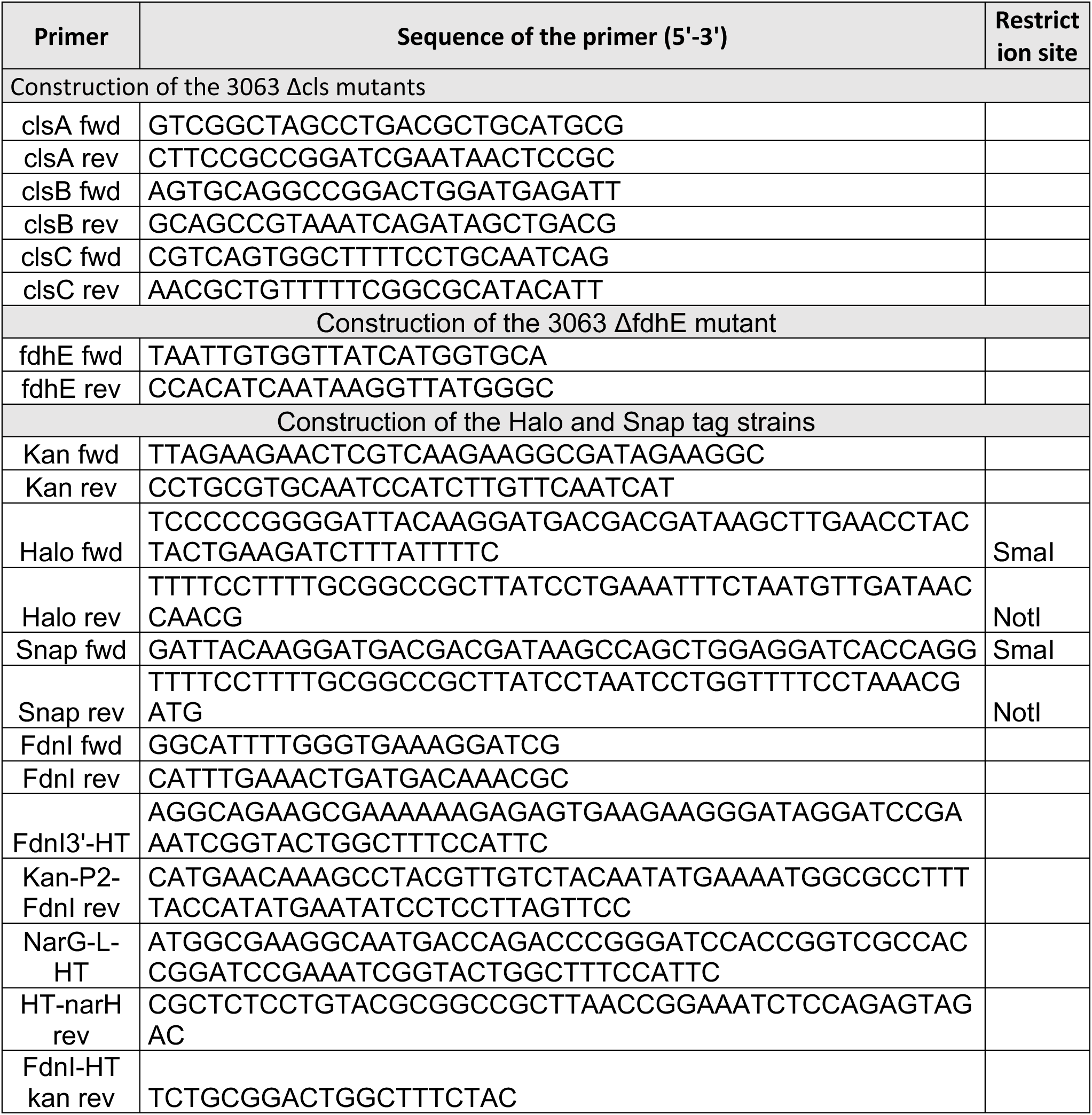
Oligonucleotides and synthetic DNA sequences used in this study.

## *SNAPtag* (Gencust)

ATGGACAAAGATTGCGAAATGAAACGTACCACCCTGGATAGCCCGCTGGGCAAACTGGAACTGAGCGGCTGCGAACAGGGCCTGCATGAAATTAAACTGCTGGGTAAAGGCACCAGCGCGGCCGATGCGGTTGAAGTTCCGGCCCCGGCCGCCGTGCTGGGTGGTCCGGAACCGCTGATGCAGGCGACCGCGTGGCTGAACGCGTATTTTCATCAGCCGGAAGCGATTGAAGAATTTCCGGTTCCGGCGCTGCATCATCCGGTGTTTCAGCAGGAGAGCTTTACCCGTCAGGTGCTGTGGAAACTGCTGAAAGTGGTTAAATTTGGCGAAGTGATTAGCTATCAGCAGCTGGCGGCCCTGGCGGGTAATCCGGCGGCCACCGCCGCCGTTAAAACCGCGCTGAGCGGTAACCCGGTGCCGATTCTGATTCCGTGCCATCGTGTGGTTAGCTCTAGCGGTGCGGTTGGCGGTTATGAAGGTGGTCTGGCGGTGAAAGAGTGGCTGCTGGCCCATGAAGGTCATCGTCTGGGTAAACCGGGTCTGGGA

## *HaloTag* (Genecust)

GGATCCGAAATCGGTACTGGCTTTCCATTCGACCCCCATTATGTGGAAGTCCTGGGCGAGCGCATGCACTACGTCGATGTTGGTCCGCGCGATGGCACCCCTGTGCTGTTCCTGCACGGTAACCCGACCTCCTCCTACGTGTGGCGCAACATCATCCCGCATGTTGCACCGACCCATCGCTGCATTGCTCCAGACCTGATCGGTATGGGCAAATCCGACAAACCAGACCTGGGTTATTTCTTCGACGACCACGTCCGCTTCATGGATGCCTTCATCGAAGCCCTGGGTCTGGAAGAGGTCGTCCTGGTCATTCACGACTGGGGCTCCGCTCTGGGTTTCCACTGGGCCAAGCGCAATCCAGAGCGCGTCAAAGGTATTGCATTTATGGAGTTCATCCGCCCTATCCCGACCTGGGACGAATGGCCAGAATTTGCCCGCGAGACCTTCCAGGCCTTCCGCACCACCGACGTCGGCCGCAAGCTGATCATCGATCAGAACGTTTTTATCGAGGGTACGCTGCCGATGGGTGTCGTCCGCCCGCTGACTGAAGTCGAGATGGACCATTACCGCGAGCCGTTCCTGAATCCTGTTGACCGCGAGCCACTGTGGCGCTTCCCAAACGAGCTGCCAATCGCCGGTGAGCCAGCGAACATCGTCGCGCTGGTCGAAGAATACATGGACTGGCTGCACCAGTCCCCTGTCCCGAAGCTGCTGTTCTGGGGCACCCCAGGCGTTCTGATCCCACCGGCCGAAGCCGCTCGCCTGGCCAAAAGCCTGCCTAACTGCAAGGCTGTGGACATCGGCCCGGGTCTGAATCTGCTGCAAGAAGACAACCCGGACCTGATCGGCAGCGAGATCGCGCGCTGGCTGTCTACTCTGGAGATTTCCGGT

